# Directional alignment of different cell types organizes planar cell polarity

**DOI:** 10.1101/2025.05.13.653899

**Authors:** Masaki Arata, Hiroshi Koyama, Toshihiko Fujimori

**Affiliations:** Division of Embryology, National Institute for Basic Biology, Okazaki, Aichi, Japan; SOKENDAI (The Graduate University for Advanced Studies), Okazaki, Aichi, Japan

## Abstract

Planar cell polarity (PCP) refers to unidirectional coordination of cell polarity in the plane of epithelial tissues, whose proper formation is essential for development in various tissues. While imbalances in the amount of core PCP proteins (core proteins) between adjacent cells disrupt PCP in some tissues, other tissues maintain PCP despite containing multiple cell types with different levels of core proteins. How such tissues tolerate these imbalances remains unclear. Here, we theoretically analyzed the contribution of spatial distribution of different cell types to PCP maintenance. We adopted a previously established PCP model on two-dimensional cell sheets, where local protein-protein interactions were assumed. Our systematic simulations revealed that the patterns of cell-type distribution significantly affect the accuracy of PCP maintenance under the condition of imbalanced PCP proteins. To identify the critical patterns, we applied both deep learning techniques and statistical modeling to the simulation data. Consequently, these analyses revealed that orientation of cell-type alignment is a key aspect of cell-type distribution that affects PCP. Such directional cell-type alignment was observed in the mouse oviduct. Our findings highlight the overlooked contribution of spatial distribution of cell types to PCP maintenance in tissues with multiple cell-types.

## Introduction

The proper function of various organs depends on planar cell polarity (PCP), the tissue-level coordination of cell polarity within the plane of epithelial tissue (1–8). For example, the transport of oocytes is driven by the ovary-to-uterus-directed beating of cilia in the oviduct epithelium (9–12), and misoriented hair cells in the cochlea are associated with hearing deficit (13).

Previous studies have identified an evolutionally conserved group of proteins, known as core PCP proteins (hereafter referred to as core proteins), that play key roles in PCP establishment (1–7). Core proteins form two types of complexes at cell edges (Fig. 1A): one comprised of CELSR and FZD (FRIZZLED), and another comprised of CELSR and VANGL (Van Gogh-like; hereafter referred to as the FZD complex and the VANGL complex, respectively). In various developing organs, the FZD and VANGL complexes show asymmetric distributions at cell edges along body axes (1, 4, 8). For example, in the mouse oviduct, the FZD complex is localized to the uterus side, while the VANGL complex is localized to the ovary side of the cell (10). This asymmetry is established before ciliogenesis, and the loss of CELSR1 results in misoriented ciliary beating (9), suggesting that the asymmetry is a prerequisite for PCP establishment. Although details remain unclear, experimental and theoretical studies suggest that the establishment of the asymmetry requires intracellular and intercellular interactions between the FZD and VANGL complexes (14– 17).

**Figure 1.**
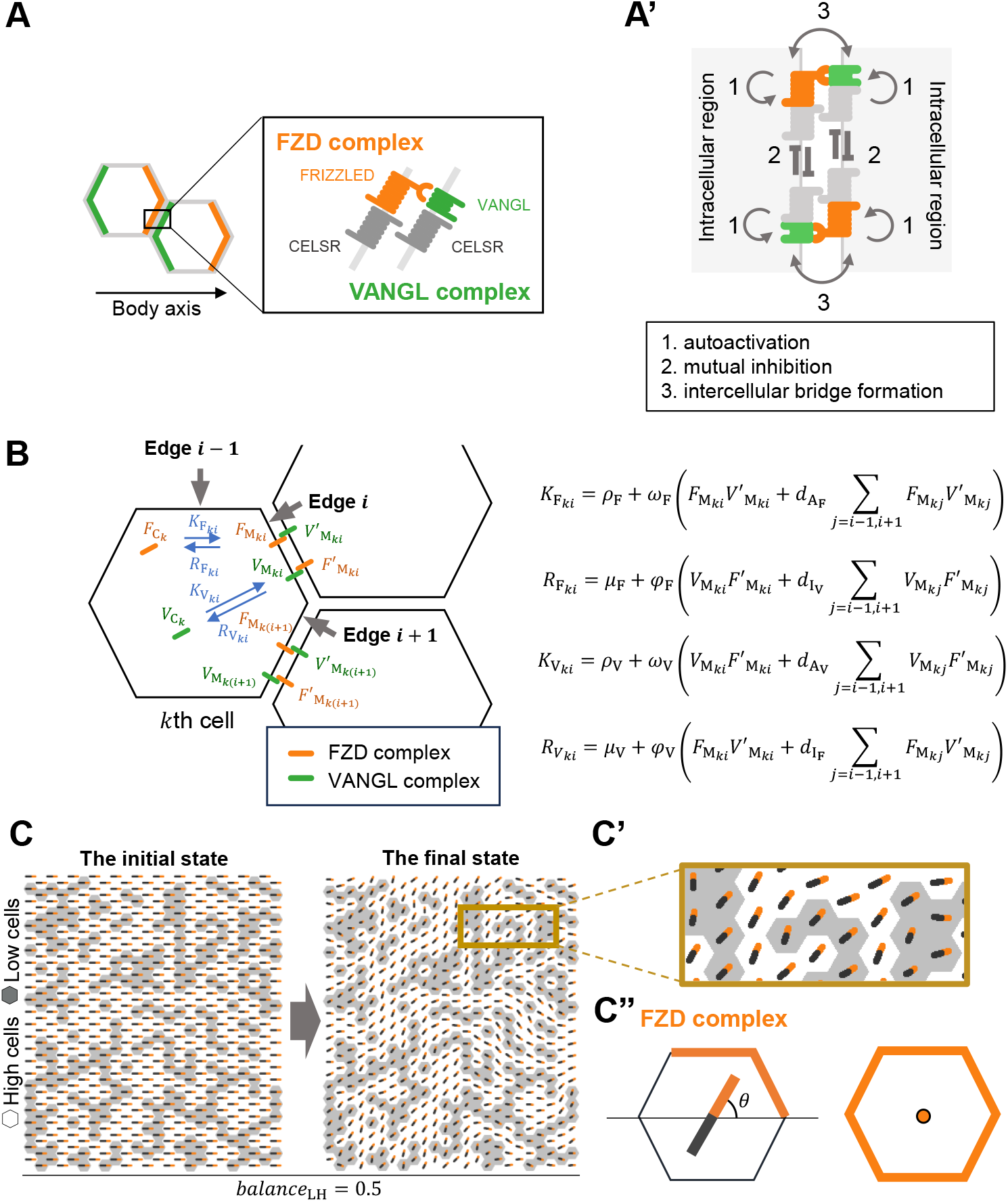
Mathematical modeling for evaluating effects of spatial organization of core protein imbalance on PCP. (**A**) A diagram illustrating the asymmetric distribution of the FZD and VANGL complexes on cell edges. (**A’**) Three key interactions between core-protein complexes implemented in our mathematical model: 1. Autoactivation: the FZD and VANG complexes stabilize complexes of the same type. 2. Mutual inhibition: the FZD and VANGL complexes mutually destabilizes complexes of the opposite type. 3. Intercellular bridge formation: the FZD and VANG complexes form bridges, intercellularly. (**B**) Diagram illustrating an overview of the mathematical model. See the main text and Supporting Information for details. (**C-C”**) Example analysis of how cell-type distribution affects PCP. White hexagons: high cells; gray hexagons: low cells. (**C**) The unidirectionally biased distribution of the FZD and VANGL complexes was reorganized by core protein imbalance (*balance*_LH_ = 0.5, compare the initial and final states). A boxed region in (**C**) is enlarged and shown in (**C’**). (**C”**) A bar centered in each cell represents polarity of cell-edge distribution of the FZD complexes. its length represents the magnitude of polarity, and its angle represents the orientation (see Supplementary Information for details).

Epithelial tissues are typically composed of multiple cell types. In such cases, how is PCP formed or maintained? The mouse oviduct epithelium consists of two cell types: multiciliated cells (MCCs) and secretory cells (SCCs, Fig. 1A). Notably, these two cell types differ in their levels of core proteins; CELSR1 is enriched at the cell edges of MCCs but is present at lower levels in SCCs (Fig 1B-B”, (9)). This imbalance in core protein levels between adjacent cells (hereafter referred to as core protein imbalance) has the potential to disrupt PCP. Indeed, when cells lacking Frizzled (Fz, a fly homolog of FZD), Van Gogh (Vang, a fly homolog of VANGLs), or Flamingo (Fmi, also known as Starry night, a fly homolog of CELSR) are experimentally produced in the *Drosophila* wing, polarities in surrounding wild-type cells are reorganized (6, 7, 18). Nevertheless, in the oviduct, both the ciliary beating orientation and core protein distribution remain biased along the ovary-uterus axis (Fig. S1B-B”). Such core protein imbalances have also been observed in various organs, including the mouse inner ear (19), airway (20), and *Drosophila* eye disc (21). However, the mechanisms by which tissues overcome core protein imbalance to maintain PCP remain unclear.

Mathematical models have been widely used to dissect the mechanisms of PCP establishment and maintenance. Several types of PCP models exist. Mechanistic models explicitly simulate the dynamics of molecules relevant to individual core proteins or their complexes (14, 15, 22–25). On the other hand, phenomenological models abstract these molecular details and represent cell polarity as a vector that evolves according to effective parameters, such as neighbor interactions, global cues, or cell geometry (26–28). These models have been used to investigate core protein functions by performing *in silico* perturbation experiments in which clusters of mutant cells lacking core proteins are embedded within a wild-type cell sheet. These studies primarily focused on elucidating the molecular functions of core proteins, rather than studying PCP establishment and maintenance in tissues of multiple cell-types with various spatial distributions. Therefore, these studies considered just idealized configurations of mutant cell clusters: a linear cell row, a rectangular cluster (14, 15, 22). Consequently, it remains largely unclear how various spatial patterns of cell types with differing core protein levels—beyond a simple mutant cluster—affect PCP in normal tissues.

In this study, to consider the core protein imbalances, we adopted a mechanistic mathematical model in which concentrations of the FZD and VANG complexes are explicitly represented. We systematically varied the spatial patterns of core protein levels in a two-dimensional cell sheet and analyzed the resulting PCP outcomes. To identify key spatial features that promote PCP maintenance, we combined simulations with deep learning, and statistical modeling. Our analysis revealed that the directional alignment of cell types, specifically, the alignment of cell types along the tissue axis, contributes to preservation of PCP in the presence of core protein imbalance. In consistent with this theoretical finding, we found that SCCs are predominantly aligned along the ovary–uterus axis in the mouse oviduct, and PCP defects emerge in regions where this alignment is disrupted. Our study offers crucial insights into how the spatial patterning of multiple cell types influences PCP establishment and maintenance.

## Results

### Modeling PCP in a 2D cell sheet

To examine how distributions of cell types with different core protein levels affect PCP, we constructed a mathematical model (see also Supporting Information), which builds upon previous frameworks (15, 23). In our model, we explicitly implemented concentrations of the FZD and VANGL complexes at cell edges and within the cytoplasmic region (Fig. 1B). This allows us to directly address the effects of intercellular imbalances in core protein levels—an aspect not captured by phenomenological vector-based models (26–28). The epithelial sheet is represented as an array of regular hexagonal cells (Fig. 1B-C”), where the FZD complex and the VANGL complex shuttle between cell edges and the cytoplasmic region (Fig. 1B, blue arrows)

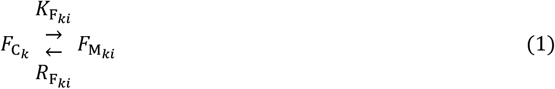

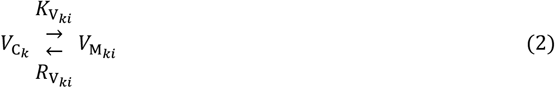

Here,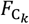and 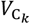 denote the concentration of the FZD and VANGL complexes in the cytoplasmic region of *k*th cell, respectively.The concentration of the FZD and VANGL complexes at *i* th cell edge in the *k*th cell are represented as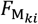and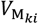, respectively; *i* = 1∼6 . In addition, concentrations of the FZD and VANGL complexes were assumed to be uniform within each cell edge and cytoplasmic region. The FZD complex shuttles between cell edge *i* and a cytoplasmic region at rates 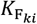and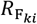, while the Vangl complex undergoes similar transport at rates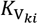and 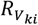. Therefore, the dynamics of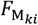 is described as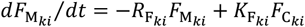; the differential equations for other components are shown in Supporting Information.

According to previous studies, to establish the planar polarized distributions of the two complexes on cell edges, the formation of intercellular bridges composed of the two complexes (the FZD-VANGL bridges) was required, where the FZD-VANGL bridges have activities of both autoactivation and mutual inhibition (Fig. 1A’; (14–17)). We basically adopted the theoretical formulation in these studies to model the above shuttling rates as follows:

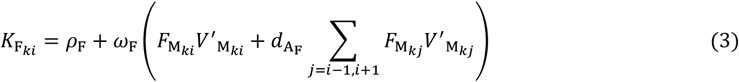

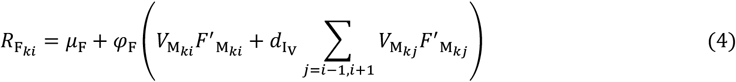

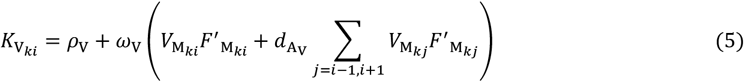

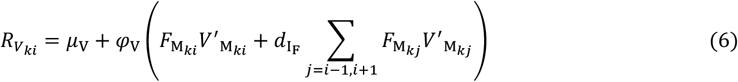

where concentrations of the FZD and VANGL complexes on a neighboring cell’s edge juxtaposing edge *i* are denoted as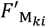 and 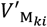,respectively (Fig. 1B). The intercellular bridges are formed between the FZD and VANGL complexes: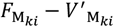and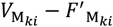. For simplicity, we assumed that the concentrations of the FZD-VANGL bridges were proportional to the terms of 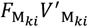 and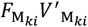 in the above equations (15). The first term in the parenthesis of eq.3 and 5 correspond to the autoactivation activity: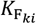and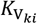 increase when 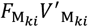and 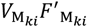are elevated, respectively. The first term in the parenthesis of eq.4 and 6 corresponds to the mutual inhibition activity: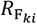 and 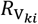increase when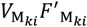 and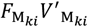 are elevated, respectively. In addition, this model assumed a non-local effect, where the shuttling rates on cell edge *i* was affected by core proteins on neighboring edges *i* − 1 and *i* + 1 (Fig. S2A and A’; the second term in the parenthesis of eq.3-6; see Supporting Information for details). A constant *d* determines the strength of the non-local effects (Fig. S2A and A’). When *d*_A_ > 0, the complexes on edge *i* − 1 and *i* + 1 elevate 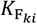and 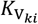. Similarly, when *d*_I_ > 0, the complexes on edge *i* − 1 and *i* + 1 elevate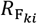 and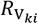 . Although the non-local effect has not been experimentally validated, theoretical studies suggest it is necessary for robust PCP establishment (15, 29).

The parameter *ρ* represents the basal internalization rate of the FZD and VANGL complexes from the cell edge to the cytoplasm, while *μ* denotes the basal trafficking rate of the FZD and VANGL complexes from the cytoplasm to the cell edge; both parameters were set to be constant. The values of parameters used in our model are listed in Table S1. We selected these values to reproduce the non-autonomous effects of mutant cells lacking core proteins, as observed *in vivo* (Fig. S3; see details in the Supporting Information).

### Setting of simulation and definition of polarity index

Before showing systematic analyses of simulations, we explain the setting of simulations. First, we prepared a cell sheet in which PCP was already established (Fig. 1C, left panel). Second, core protein levels were reduced in selected cells (low cells; gray cells in Fig. 1C; remaining cells are referred to as high cells in the following). The magnitude of the core protein imbalance was defined as a constant *balance*_LH_, which ranges from 0 to 1. When *balance*_LH_ = 1, low and high cells contain equal amounts of core proteins, whereas, when *balance*_LH_ = 0, low cells have no core proteins. Then, we performed simulations to analyze how the distribution of core proteins was reorganized. Fig. 1C is an example in the case that low cells (*balance*_LH_ = 0.5) are randomly distributed. At the initial state, distributions of core proteins at cell edges were unidirectionally biased, with the FZD and VANGL complexes enriched at right and left ends of each cell, respectively (Fig. 1C and S2B). In our data, we represent each cell’s polarity vector, which indicates an orientation of FZD complex-enriched cell edges, with an orange-headed bar (Fig. 1C”, see details in Supporting Information). The bar’s length corresponds to the polarity magnitude, and its angle corresponds to the polarity direction.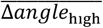is the mean absolute change in polarity vector angles of high cells in a cell sheet. Since the value of 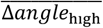 is 0 at the initial state of the simulation, the value of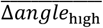 at the final state represents the extent of polarity reorganization. In Fig. 1C,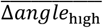 is 36.0 degrees at the final state, indicating that the core protein imbalance disrupts PCP.

### Systematic simulations under various cell type distribution

To gain insights into which aspects of cell-type distributions influence PCP, we generated cell-type distributions by randomly placing low cells, expecting these distributions to encompass diverse features of cell-type distributions. Through our simulations, we evaluated their effects on PCP by calculating 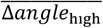 and searched for features of cell-type distributions that significantly impact PCP.

We began by generating random distributions with low-cell frequencies (*freq*_low_) ranging from 0.05 to 0.95 in increments of 0.05. At *freq*_low_ = 0, all cells are high cells, whereas at *freq*_low_ = 1, all cells are low cells. For each value of *freq*_low_, we generated 10 random distributions and performed simulations (Fig. 2A-B). Fig. 2A-A” shows examples of our simulations at *balance*_LH_ = 0.5. *freq*_low_ significantly influenced the values of 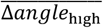(Fig. 2B; see the boxed column for these values at *balance*_LH_ = 0.5), indicating that *freq*_low_ is an essential parameter influencing PCP. Additionally, changes in *balance*_LH_ significantly altered the value of 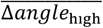(Fig. 2B and S4A-B’; e.g., compare Fig S4A with S4A’). We further noted that the influence of low cells on 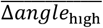 was particularly evident when *balance*_LH_ was below 0.6 (Fig. 2B). This indicates that PCP is affected by low cells even when core proteins are only partially lost in these cells.

**Figure 2.**
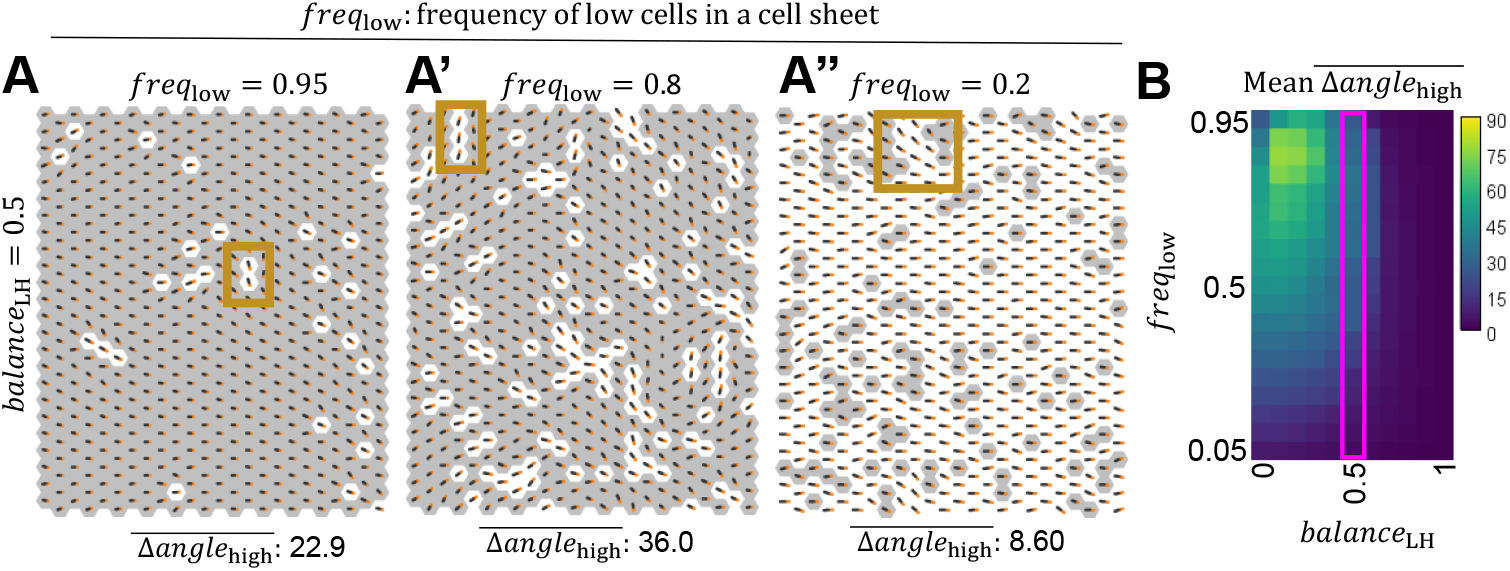
Systematic evaluation of effects of random low-cell distributions and magnitude of core protein imbalance on PCP. (**A-A”**) Examples showing the effects of random low-cell distributions on PCP. *balance*_LH_ = 0.5. *freq*_low_ = 0.95 (**A**), 0.8 (**A’**), and 0.95 (**A”**). The resulting values of 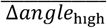 are shown at the bottom of each panel. Boxes in **A** and **A’** highlight alignment of high cells along the Y-axis of the cell sheet. A box in **A”** highlight the misoriented polarity vectors of high cells in the vicinity of low cell clusters. (**B**) For each *freq*_low_ value, 10 random distributions of low cells were generated and their effects on PCP were evaluated by simulation. Heatmap colors represent the average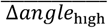values over 10 random distributions for each combination of *balance*_LH_ and *freq*_low_ (unit: degrees). The boxed column represents data at *balance*_LH_ = 0.5.

Although *freq*_low_ and *balance*_LH_ appeared to be essential parameters influencing 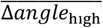, they might not be the only determinants of 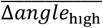 . For example, among the 10 simulations under *freq*_low_ = 0.4 and *balance*_LH_ = 0.5 with different cell-type distributions, the maximum and the minimum values of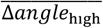 were 36.4 and 16.3 degrees, respectively (variance: 40.5 degrees; Fig. S4B). Variances of 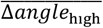 at different values of *freq* and *balance* are summarized in Fig. S4C. We expected that such a variance can be explained by differences in features of cell-type distribution between cell sheets. One candidate is the orientation of the alignment of low and high cells. As highlighted by boxes in Fig. 2A-A’, polarity vectors of high cells were strongly reoriented where these cells were aligned along the Y-axis of the cell sheets. In addition, we noticed that when high cells were dominant in a cell sheet, polarity vectors of high cells were reoriented only in the vicinity of low cells (see a boxed region in Fig. 2A”), suggesting that differences in the frequency of edges between high and low cells underlie the variations in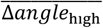 between cell sheets. This is consistent with a previous theoretical study, which report that that core proteins were eliminated from cell edges between Fmi (Flamingo, a fly homolog of CELSRs)-deficient cells and wild type cells (22). We analyzed effects of these parameters on the values of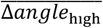 in later sections.

### Cell type distribution is effective on PCP maintenance

To examine whether features of cell-type distribution can explain differences in values of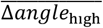 between cell sheets, we set out to train a neural network to predict the extent of PCP disorganization in a cell sheet using only information on cell-type distributions (Fig. 3A). If the network was successfully trained, it indicates that the cell-type distribution is a crucial factor in explaining the value of 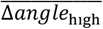 in a cell sheet. Otherwise, it implies that the cell-type distribution has little impact.

**Figure 3.**
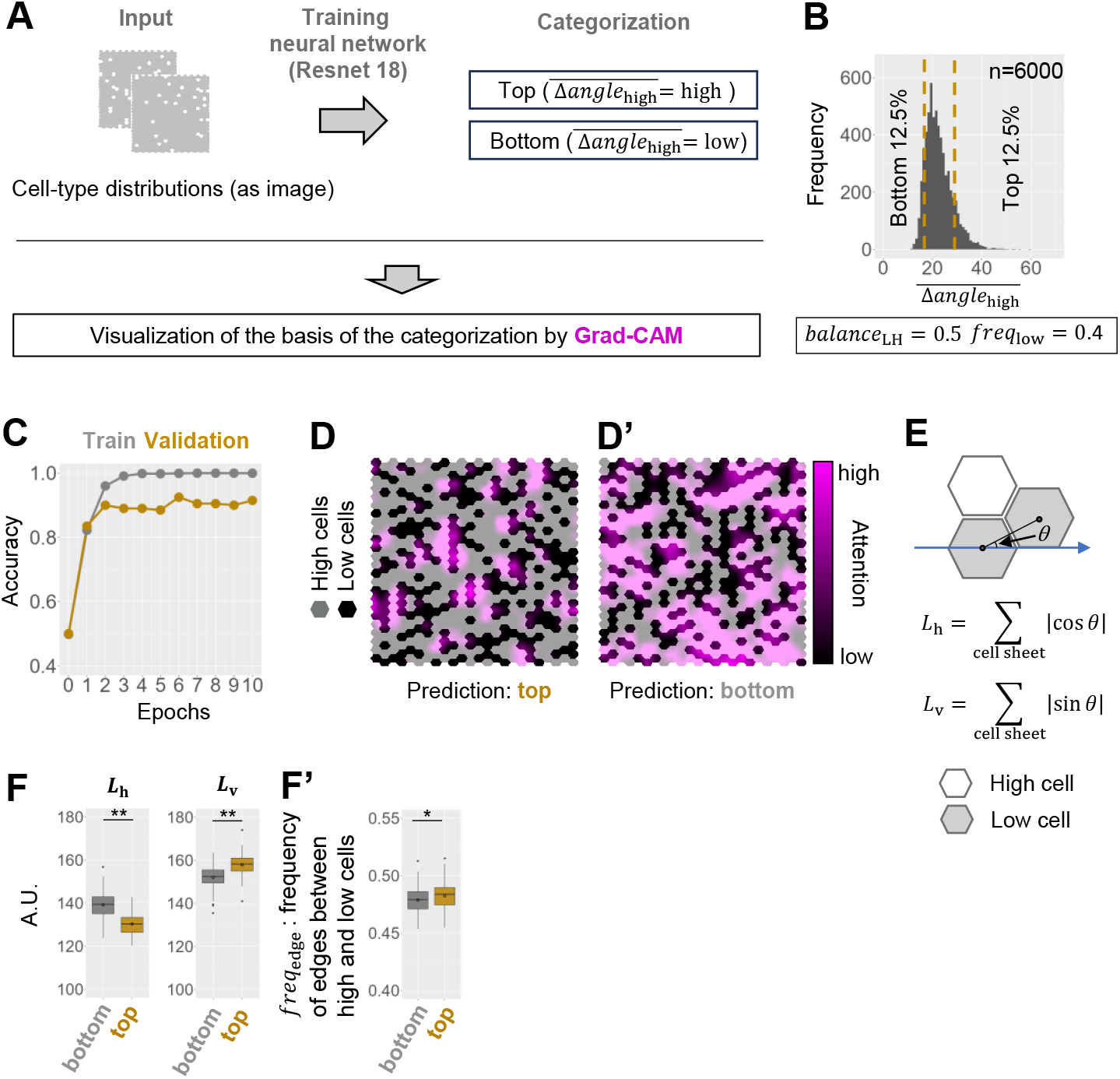
Exploring features of cell-type distribution affecting PCP by deep learning techniques. (**A**) Diagram illustrating the strategy for deep learning analysis. Cell-type distributions were provided as images for the input into the neural network. No information about polarity vectors was included in the input images. The neural network (ResNet18) was trained to predict whether a given cell-type distribution corresponds to a “top”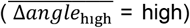 or “bottom”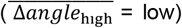 classification. Finally, Grad-CAM was applied to visualize the regions in the input images that contributed to the prediction. (**B**) A total of 6,000 random distributions were generated at *freq*_low_ = 0.4, and their effects on PCP were simulated at *balance*_LH_ = 0.5. The variation in the resulting 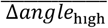 values is shown as a histogram. Distributions within the top 12.5% and bottom 12.5%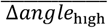values were labeled “top” and “bottom”, respectively. (**C**) Prediction accuracy of the neural network during training. The gray and gold lines show changes in prediction accuracy for the training and validation data, respectively. (**D-D’**) Grad-CAM attention maps highlighting regions of input distributions where the network emphasized during predictions. Black and gray hexagons represent low and high cells, respectively. Magenta brightness indicates attention magnitude. (**D**) A distribution predicted as “top”. Attention is concentrated at vertical edges between low cells and high cells. (**D’**) A distribution predicted as “bottom”. Attention is concentrated around low cells adjacent to high cells. (**E**) Diagram illustrating the definition of *L*_h_ and *L*_v_. Gray and white cells represent low and high cells, respectively. Blue arrow indicates the X-axis of the cell sheet. (**F**) Boxplots showing *L*_h_ and *L*_v_ values for top and bottom validation data (**: p<0.01; Wilcoxon rank-sum test). (**F’**) Boxplots showing frequencies of edges between low cells and high cells for the top and bottom validation data (*: p<0.05; Wilcoxon rank-sum test).

To this end, we first generated a large number of simulation data with *freq*_low_ = 0.4 and *balance*_LH_ = 0.5, as we anticipated that this parameter set would produce a wide range of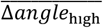values. A total of 6,000 random cell-type distributions were generated, and those in the top 12.5% and bottom 12.5% of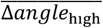 were labeled as “top” and “bottom” distributions, respectively (Fig. 3A and B). In addition to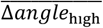, we calculated the averaged angle of polarity vectors over low cells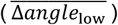and all cells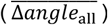. However, using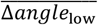or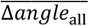for labeling did not significantly alter the classification of each distribution as “top” or “bottom”, since these values were closely correlated with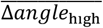 (Fig. S4D and E).

Then, a ResNet18-based neural network (30) was trained to categorize distributions as either “top” or “bottom” (Fig. 3A). Among the 750 “top” distributions and 750 “bottom” distributions, 100 from each group were used for validation, while the remaining distributions were used for training. After training, the model achieved a classification accuracy of 0.925 (Fig. 3C), indicating that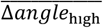 is determined by cell-type distributions.

### Deep learning-based exploration of features of cell type distributions that affect PCP

To find what kind of spatial cell-type distributions is critical for PCP maintenance, we used an explainable artificial intelligence technique, Grad-CAM (31), by which we can visualize regions of a cell sheet where the network focused its attention to make predictions (Fig. 3A). We expected that this technique would allow us to identify key features of cell-type distributions that influence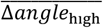. Notably, in distributions predicted as “top,” Grad-CAM highlighted regions where low cells were aligned along the Y-axis of the cell sheet (Fig. 3D). In contrast, in distributions predicted as “bottom,” Grad-CAM highlighted regions where low cells were aligned along the X-axis of the cell sheet (Fig. 3D’). To quantify these patterns, we measured the total lengths of horizontal and vertical alignment lengths of low cells adjacent to high cells (Fig. 3E). For each pair of adjacent low cells whose shared vertex was also shared by a high cell, we measured the alignment angle *θ* between the two low cells (Fig. 3E). Here, *θ* is defined as the angle between the line connecting the centroids of the two low cells and the X-axis of the cell sheet (Fig. 3E). We then calculated the total low-cell alignment as follows:

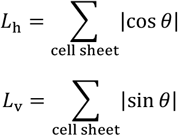

where *L*_h_ and *L*_v_ are total horizontal and vertical alignment lengths of low cells adjacent to high cells, respectively. Hereafter, we refer to alignment of low cells adjacent to high cells simply as “low-cell alignment.”

The values of vertical alignment (*L*_v_) of “top” distributions were significantly higher than those of “bottom” distributions (p<0.01; Student’s *t*-test; Fig. 3F). In contrast, the values of horizontal alignment (*L*_H_) of “top” distributions were significantly lower than those of “bottom” distributions (p<0.01; Student’s *t*-test; Fig. 3F). As discussed earlier, we expected that the frequency of edges between high cells and low cells (*freq*_edge_) would influence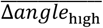. Although the difference was subtle, we found that the value of *freq*_edge_ was higher in “top” distributions than in “bottom” distributions (p<0.05; Student’s *t*-test; Fig. 3F’). As discussed later, the contribution of this parameter to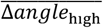 is not negligible.

### GLM-based evaluation of factors affecting PCP

Finally, to dissect effects of orientations of low-cell alignment and other aspects of cell-type distributions on PCP, we constructed a generalized linear model (GLM), a framework for modeling the relationship between an object variable and explanatory variables. Here, the object variable was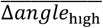 and candidates for explanatory variables included *L*_h_, L_v_, b*alance*_*L*H_, *freq*_low_, and *freq*edge . In short, by using GLM, we can expect to evaluate to what extent each explanatory variable contributes to the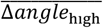. Based on the value of AIC (Akaike Information Criterion), we selected the model that best predicted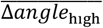in our simulation data (Fig. 4). Data with *balance*_LH_ >0.6 were excluded from the GLM analysis, as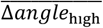 was weakly affected by cell distributions when the *balance*_LH_ was high. In addition, since *L*_h_ and *L*_v_ were strongly correlated (i.e., positive correlation) in the dataset, their direct inclusion in the model was problematic. To address this, we converted them into a single variable, *L*_h/v_ = L_h_/*L*_v_, which represents the anisotropy of low-cell alignment. The optimal model selected based on the value of AIC was as follows:

**Figure 4.**
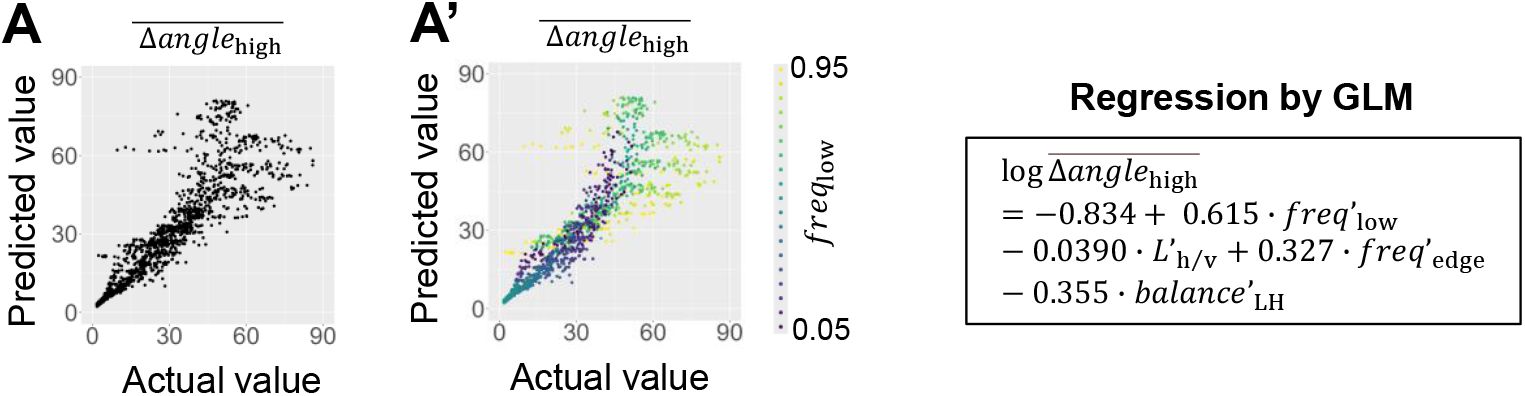
Regression analysis of features of cell-type distributions by Generalized Linear Model (GLM). (**A-A’**) Scatter plots showing the correlation between actual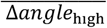values and those predicted by the GLM. Each dot represents a data point shown in Figure 2**B.** *freq*_low_ values are color-coded in (**A’**). The detail of the model is shown in the right panel. Pearson correlation coefficient: 0.859.

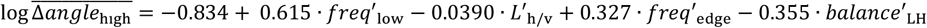

where values of explanatory variables were normalized so that their mean and SD were 0 and 1, respectively. Normalized variables are marked with an apostrophe (e.g.,*freq*′_*low*_). The values of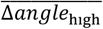 predicted by this model were closely correlated with actual values of 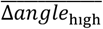 (Fig. 4A; Pearson correlation coefficient: 0.859). However, the model’s accuracy decreased when the frequency of low cells was high, particularly at *freq*_low_= 0.95 (yellow dots in Fig. 4A’). Here, the absolute value of each coefficients indicates the strength of the influence of explanatory variables on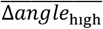 . The coefficients indicate that low cell frequencies (*freq*_low_) and magnitude of core protein imbalances (*balance*_LH_) have a significant impact on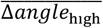, as expected. In addition, the model suggests that the frequency of edges between low and high cells (*freq*_edge_) is a key parameter. Although the absolute value of the coefficient for the anisotropy of low-cell alignment (*L*^′^_h/v_) was smaller than those of the other explanatory variables, a model excluding *L*_h/v_ was less optimal according to AIC. Since the coefficient of *L*_h/v_ is negative, the model suggests that an increase in the ratio of total lengths of horizontal to vertical alignment of low cells decreases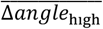. These results indicate that the frequency of low cells (*freq*_low_), the frequency of edges between high and low cells (*freq*_*edge*_), and the anisotropy of low-cell alignment (*L*_h/v_) are key aspects of cell-type distributions that influence PCP.

### The direction of low-cell alignment is essential for mitigating the effect of low cells on PCP

To further address whether the direction of low-cell alignment affects PCP, we placed rectangular low-cell clusters within a cell sheet and analyzed the degree of PCP reorganization after simulations. First, low-cell clusters were aligned in a lattice at intervals of 3 cells and *balance*_LH_ was set to 0 (i.e., no core proteins in the low cells). We then examined the effects of low-cell clusters with varying values of *l*_h_ and *l*_v_ (Fig. 5A-A’). Here, *l*_h_ and *l*_v_ represent the lengths of the cluster in the horizontal and vertical directions, respectively, relative to the X-axis, which corresponds to the initial direction of PCP. Our analysis revealed that when low-cell clusters were elongated perpendicular to the X-axis, the value of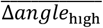 increased (Fig. 5A, compare *l*_h_ = 3, *l*_v_ = 5 with *l*_h_ = 3, *l*_v_ = 3). In contrast, when low-cell clusters were elongated parallel to the X-axis, the values of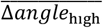 did not increase (Fig. 5A, compare *l*_h_ = 5, *l*_v_ = 3 with *l*_h_ = 3, *l*_v_ = 3).

**Figure 5.**
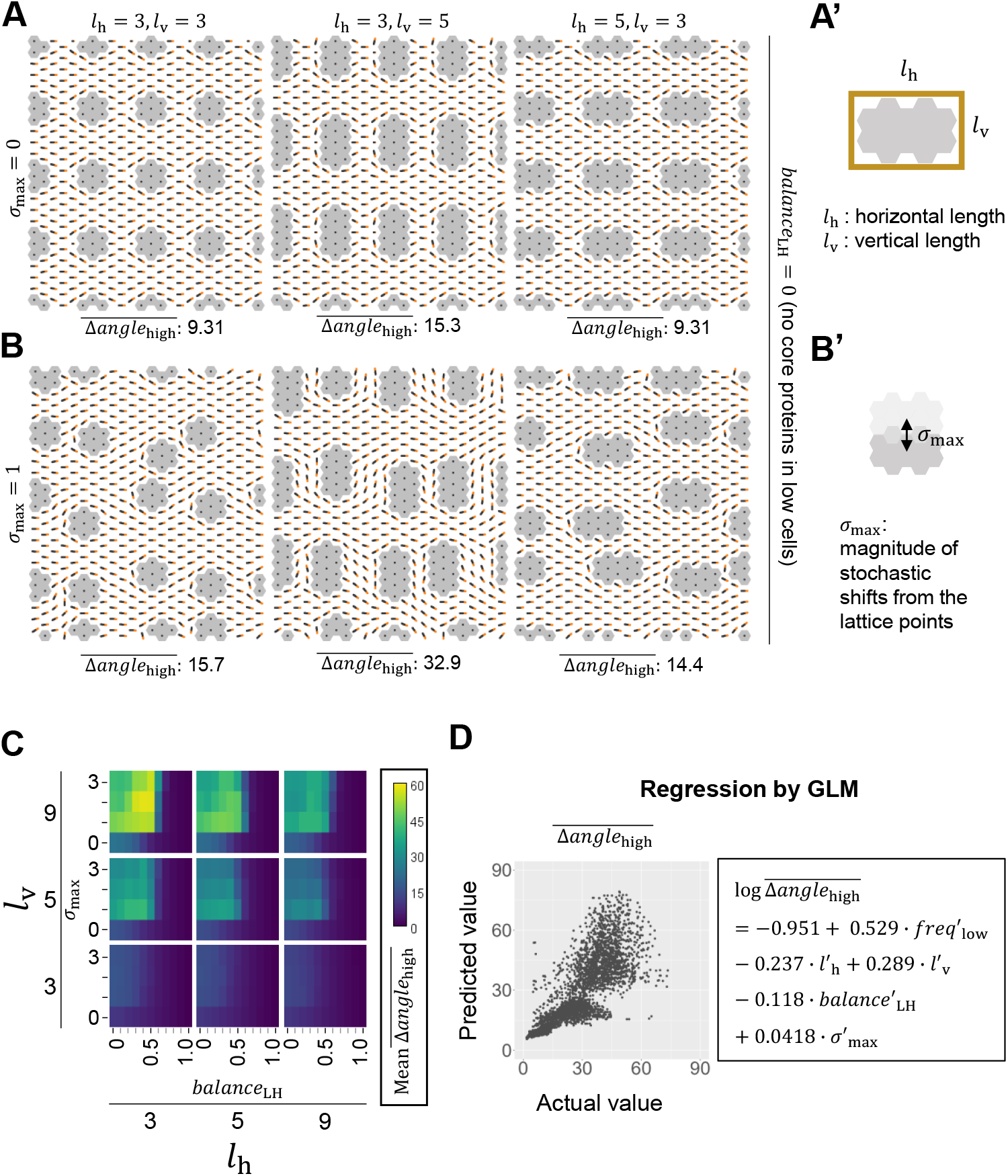
Evaluating the effects of directions of low-cell alignment on PCP. (**A-B’**) Examples of simulations that evaluate the effects of low-cell cluster lengths (**A’**) on PCP. The resulting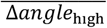 values are shown at the bottom of each panel. Clusters were arranged in a lattice (**A**) and were randomly shifted up to σ_max_cell(s) along the X and Y axis of each cell sheet (**B** and **B’**). (**C**) Heatmaps showing mean values of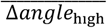, averaged over 10 distributions for each combination of *balance*_LH_, σ_max_, *l*_h_, and *l*_v_ . Mean values of 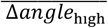 are color coded as shown in the right lookup table (unit: degrees). (**D**) Scatter plot showing a correlation between actual values of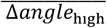 and those predicted by the GLM (right panel). Pearson correlation coefficient: 0.822. Each dot represents a single simulation.

Next, we introduced stochastic shifts from the lattice points (Fig. 5B-B’). The parameter σ_max_ determines the magnitude of this shift. When σ_max_= 1, low cell clusters could be displaced by up to one cell along both the x- and y-axis. Fig. 3B shows that elongation of low-cell clusters along the Y-axis increased the mean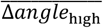, while elongation of low-cell clusters along the X-axis did not (*l*_h_ = 3, l_v_ = 5 vs. *l*_h_ = 5, *l*_v_ = 3). We obtained similar results under the conditions of various parameter values, σ_max_(1∼3), *l*_h_, *l*_v_, and *balance*_LH_ (Fig. 5C). When low cell clusters were regularly aligned, misorientation of polarity vectors was observed only in the immediate vicinity of low cell clusters (Fig. 5A). In contrast, when clusters were shifted, misorientation of polarity vectors was not limited to the vicinity of low cell clusters (Fig. 5B, middle panel), suggesting that shifting positions of clusters disrupts PCP.

These findings support our hypothesis that the direction of low-cell alignment plays a key role in mitigating the disruptive effects of low cells on PCP. However, in our analysis, modifying the lengths of low-cell clusters inevitably altered other aspects of cell-type distributions. For instance, changes in *l*_h_ and *l*_v_ also affected the frequency of low cells. To dissect effects of low-cell alignment and other aspects of cell-type distributions on PCP, we constructed a GLM. Here, the object variable was 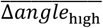 and candidates for explanatory variables included *balance*_*L*H_, the frequency of low cells (*freq*_low_), σ_max_, *l*_h_, and *l*_v_. Based on the value of AIC, we selected the model that best predicted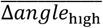 in our simulation data as follows.

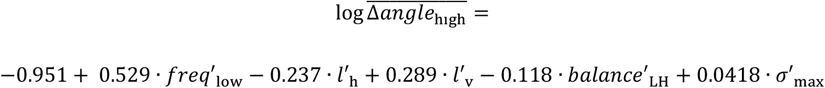

where all explanatory variables are normalized. The Pearson correlation coefficient between the actual and predicted values of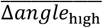 was 0.822 (Fig. 5D), indicating that the model accurately described the relationship between the object variable and explanatory variables. Among all coefficients, the absolute value of the coefficient of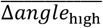is the largest, indicating that frequency of low cells has the most significant effect on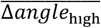 . The model shows that *l*_h_ and *l*_v_ are also essential parameters that affect PCP, as their coefficients have the second and third highest absolute values, respectively. As expected, the model shows that an increase in horizontal length (*l*_h_) reduces 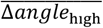, an increase in vertical length (*l*_v_) elevates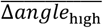 . In comparison, the absolute value of the coefficient for σ^′^_max_is much smaller than those of *l*^′^_h_ and *l*^′^_v_. Therefore, the effect of cluster distribution (σ_max_) on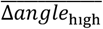 is less significant than that of cluster lengths (*l*_h_ and *l*_v_). These results were conserved in a different type of cell array (Fig. S2B’ and S5). Taken together, we concluded that the directional low-cell alignment is a key aspect of cell-type distributions that mitigates the effects of low cells on PCP.

### Orientation of PCP is correlated with the direction of low-cell alignment *in vivo*

To evaluate the relevance of our theoretical findings to living systems, we analyzed the mouse oviduct epithelium, which consists of only two cell types: secretory cells (SCCs) and multiciliated cells (MCCs). Because the frequency and distributions of SCCs are varied across different regions in the oviduct, we simultaneously analyzed both the direction of PCP and the distribution patterns of SCCs. We focused on two regions: infundibulum and ampulla.

In the infundibulum of the adult mouse oviduct, previous studies reported that SCCs comprise 24% (*freq*_low_ = 24%) of the epithelial cells (11 weeks old; 1, 3), where distributions of core proteins are biased along the ovary-uterus axis, and that CELSR1 levels are lower in SCCs than in MCCs (Fig. S1B-B”, 1). First, we quantitatively evaluated the imbalance in CELSR1 levels between SCCs and MCCs. To this end, we measured the intensities of CELSR1 immunofluorescent signals at cell edges. We found that CELSR1 signals at edges between MCCs were approximately twice as high as those at edges between SCCs (Fig. S6A-C’, *balance*_LH_ = 0.5). We also validated this conclusion by using transgenic mice expressing EGFP-fused VANGL2 ubiquitously (Fig. S6D-E’). These observations suggest that SCCs have fewer levels of core proteins than MCCs and do not establish the polarized distribution of core proteins at cell edges.

Next, we analyzed the distribution of SCCs in the infundibulum. For this purpose, we used previously-published images in which cell edges and MCCs were fluorescently labeled (Fig. 6A and A’; images were adopted from (11)). In the oviduct epithelium, SCCs form small clusters and these clusters scatter around the epithelium (Fig. 6A and A’). We measured the length of each SCC cluster by placing a bounding rectangle whose horizontal edges were aligned parallel to the ovary-uterus axis (Fig. 6A’). The mean *l*_h_ was 2.45 cells (Standard deviation, SD: 1.16), which was significantly larger than the mean *l*_v_, 1.85 cells (SD: 0.862; p<0.01, Wilcoxon rank sum test, Fig. 6B and B’). This result indicates that SCCs preferentially align along the direction of PCP in the infundibulum.

**Figure 6.**
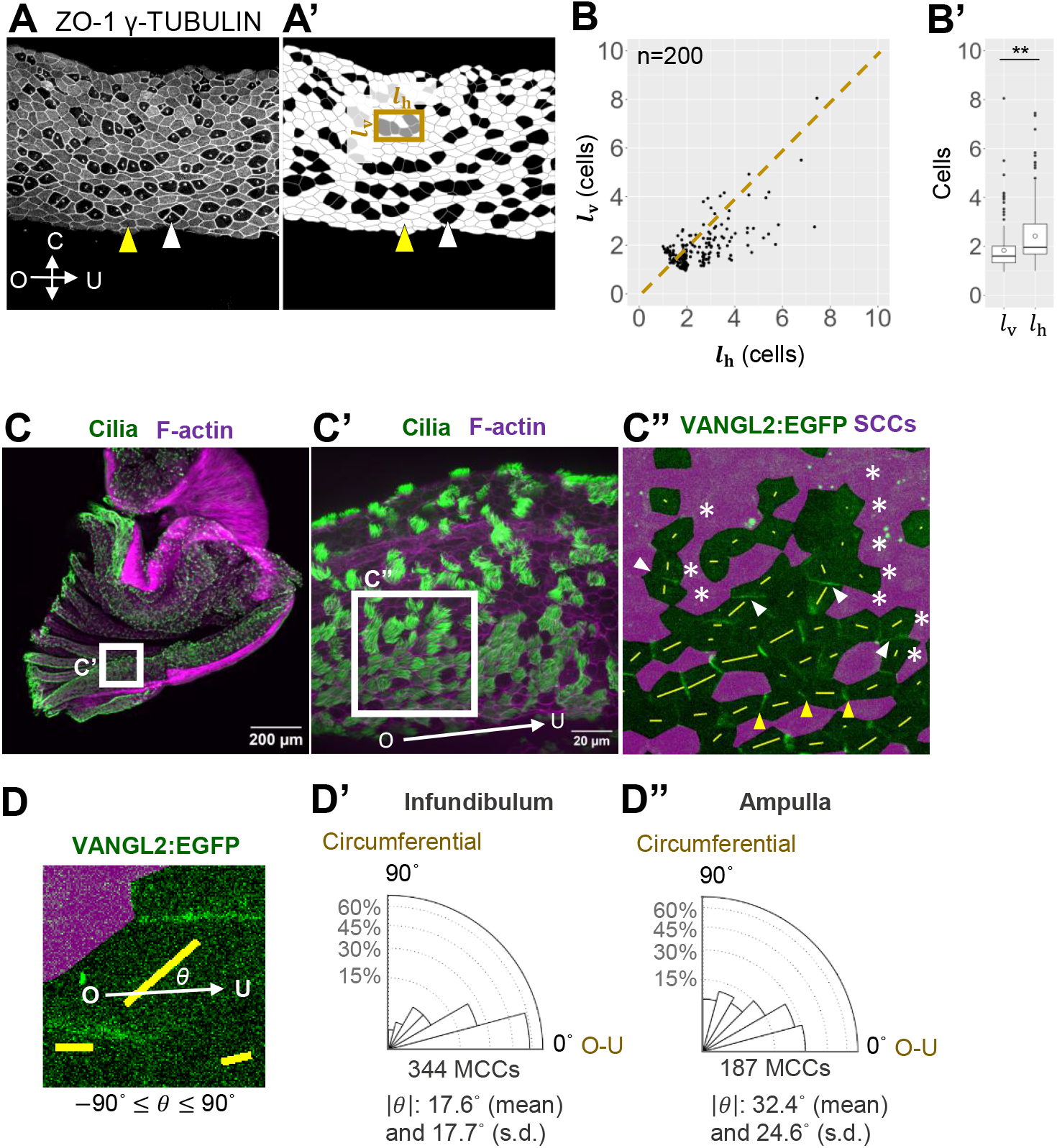
Direction of alignment of secretory cells was correlated with the orientation of core protein distribution. (**A-B’**) Analysis of SCC distribution in the infundibulum region of the oviduct. (**A**) Oviduct stained for ZO1 (a marker for cell edges) and γ-TUBULIN (a marker for MCCs). White and yellow arrowheads point to a SCC and MCC, respectively. (**A’**) Segmented image: gray lines denote cell edges; white cells are MCCs; black cells are SCCs. The ovary-uterus (O-U) and circumferential (C) lengths of each SCC cluster (*l*_h_ and *l*_v_, respectively) were measured using a bounding rectangle (gold box). (**B-B’**) *l*_h_ and *l*_v_ values were analyzed using a scatter plot (**B**) and a boxplot (**B’**). The dotted line in (**B**) represents *l*_h_ = l_v_ . Open circles in **B’** indicate average values. **: p<0.01 (Wilcoxon signed-rank test). Isolated SCCs were excluded from the analysis. N: 200 clusters; 3 mice. (**C-C”**) The ampulla region of the oviduct was opened longitudinally and stained for acetylated-TUBULIN (green, a marker for cilia) and F-actin (magenta, a marker for cell edges). The boxed regions in **C** and **C’** are shown in **C’** and **C”** respectively. The direction of the ovary-uterus axis is indicated by an arrow in **C’**. (**C”**) SCCs are filled with magenta. VANGL2:EGFP signals are shown in green. Asterisks mark examples of circumferential SCC alignment. White arrowheads mark VANGL2:EGFP signals enriched at cell edges running along the ovary-uterus axis. These enrichments were observed in MCCs adjacent to SCCs aligned circumferentially. In the same view field, enrichments of VANGL2:EGFP signals at cell edges running circumferentially were also observed (yellow arrowheads). These enrichments were found near low cells aligned along the ovary-uterus axis (e.g., compare yellow arrowheads with white arrowheads in **C”**). Yellow bars in C” represent polarity of VANGL2:EGFP distributions at cell edges. The length and angle of each bar indicate the magnitude and direction of VANGL2:EGFP polarity, respectively (see Supplementary Information for details). Compare the orientations of bars near the white arrowheads with those near the yellow arrowheads (other examples are shown in Figure S7). (**D-D”**) Quantification of VANGL2:EGFP polarity in the infundibulum (**D’**) and the ampulla (**D”**). Bar angles relative to the O-U axis (**D**) were calculated and plotted as rose diagrams (circular histograms).

In the ampulla, another region of the oviduct, MCCs are sparsely distributed (Fig. 6C-C”). Although the distribution of SCCs was varied, we sometimes found regions where SCCs were circumferentially aligned. In these regions, we analyzed the distribution of VANGL2:EGFP in MCCs adjacent to these SCCs, and found that VANGL2:EGFP was enriched at cell edges running along the ovary-uterus axis in adjacent MCCs (white arrowheads in Fig. 6C”). Note that, in regions where SCCs were aligned along the ovary-uterus axis, VANGL2:EGFP in MCCs was enriched at cell edges oriented circumferentially (yellow arrowheads in Fig. 6C”). Quantitative analysis confirmed that VANGL2:EGFP localization in the ampulla was misaligned relative to that in the infundibulum (Fig. 6D-D”). Together with the results in the infundibulum, the direction of PCP in the oviduct was consistent with our theoretical outcomes.

## Discussion

In the present study, we theoretically investigated how PCP is maintained in tissues comprising multiple cell types, one of which expresses lower level of core PCP proteins, called low cell. Our systematic simulations suggested that spatial distribution of the low cells, as well as their core protein expression levels and their population, affects proper PCP maintenance. Moreover, deep learning–based analyses using Grad-CAM revealed the impact of spatial distribution: PCP is less affected when low cells are aligned parallel to the tissue polarity. Finally, through generalized linear modeling (GLM), we successfully quantified the contribution of spatial distribution to PCP maintenance. Our results showed that the alignment of low cells has significant impact on PCP maintenance, similar to that of expression levels and population size. From these results, we propose that the spatial distribution of the low cells can be an essential factor for maintaining polarity of tissues composed of multiple cell types. Consistent with our theoretical findings, PCP orientations in distinct regions of the mouse oviduct varied in accordance with the alignment of low cells.

The establishment of PCP requires interplays of core proteins between adjacent cells. Previous studies showed that experimentally induced low cells disrupt polarity of surrounding wild-type cells (18). Nevertheless, several tissues naturally exhibit imbalances in core protein levels between cells, while still maintaining PCP. Prior theoretical work has examined how mutant cells lacking core proteins influence neighboring wild-type cells, focusing primarily on the molecular mechanisms of PCP establishment. These investigations typically employed simplified spatial configurations, a linear cell row, or a rectangular cluster (14, 15, 22), without explicitly analyzing the effect of cell-type distribution. Consequently, the effect of complex, physiologically relevant spatial arrangements on PCP maintenance has remained largely unexplored. Our theoretical analysis revealed that the impact of low cells on PCP is strongly influenced by their spatial distribution. This suggests that spatial configuration acts as a regulatory factor in PCP maintenance, functioning cooperatively with molecular interactions between core proteins. In other words, cell-type distribution itself can be considered an integral component of the PCP formation system. In *Celsr1* mutant epithelium, anisotropy of SCC alignment was lost (Fig. S7), suggesting that CELSR1 is required for proper low-cell alignment. If CELSR1 directory regulates SCC alignment, there might be a feedback regulation between core proteins and cell-type distributions. However, the mechanisms governing SCC alignment along the tissue axis remain unclear.

To examine how the degree of core protein imbalance influences PCP, we introduced a parameter, *balance*_LH_, which quantifies the relative level of core proteins in low versus high cells. This allowed us to move beyond binary mutant/wild-type modeling and analyze graded imbalances in protein levels. While our study focused on a biologically relevant case in which both FZD and VANGL complex levels are equally reduced in low cells, our model framework is flexible and can accommodate other combinations of imbalance, for example, unequal reductions in the two complexes or mixed populations of different low-cell types that might be found *in vivo*. A full exploration of these parameter spaces would require extensive simulations and is beyond the scope of the current study. However, our approach can be readily extended to such scenarios in future work.

Because our mathematical model is deterministic, the outcome of each simulation is determined by the initial configuration of cell types. This feature allowed us to apply deep learning techniques to systematically identify spatial patterns that influence PCP maintenance. Specifically, we used a neural network classifier to screen diverse configurations, and subsequently employed generalized linear modeling (GLM) to evaluate the contributions of interpretable spatial features, such as alignment of low cells, to polarity outcomes. Although the main focus of this study is the relationship between cell-type distribution and PCP maintenance, this hybrid computational framework may be broadly applicable to other problems in tissue patterning. For example, it could be adapted to investigate the effects of spatial heterogeneity in signaling molecule levels beyond core proteins, offering a general strategy for bridging complex spatial input patterns with emergent tissue-level patterning.

While our study focused on cell-type distribution, we do not exclude the possibility that additional mechanisms help mitigate the effects of core protein imbalance on PCP. For example, in the mouse airway, the frequency of MCCs (0.37 on average; (32)) is much lower than in the oviduct (0.76 on average; (9)). If non-ciliated cells in the airway are low cells, our model suggests that PCP maintenance would not be feasible in the airway epithelium. Several studies have reported that extrinsic mechanical cues, such as fluid flow can align cilia beating orientation in the ventricle (33), the airway (34), and the *Xenopus* skin (35). These mechanisms might also play a role in compensating for core protein imbalance (36).

## Materials and Methods

### Simulation

We used Octave (36) for numerical simulations and visualized the data by using Fiji (37). The epithelial tissue was modeled as an array of regular hexagonal cells, with a width and height of 24 cells each. The boundary conditions were periodic in all directions. See Supporting Information for details on our mathematical modeling and simulation.

### Exploring features of cell-type distribution that affect PCP by deep learning

#### 1. Data generation and labeling

For generating random distribution of low cells in Fig. 2, random numbers that range from 0 to 1 were assigned to each cell, and cells with top 100\**freq*_low_ % values were defined as low cells. Threshold values of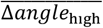for labeling “top” or “bottom” were determined by the accuracy of prediction made by the trained network. More specifically, at first, 3,000 distributions were generated at *freq*_low_ = 0.4 and *balance*_LH_ = 0.5 and data that took top 25% and bottom 25% values of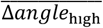 were labeled as “top” and “bottom”, respectively. We used these data for training the network, but the accuracy of the prediction made by the network did not exceed 0.85. We additionally generated 3,000 distributions and selected distributions that took top 12.5% and bottom 12.5% values of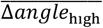 for deep learning.

Data augmentation is a technique to increase the number and diversity of training data. Since our simulation adopted the periodic edge condition in all directions, shifting a cell-type distribution along the X and Y axis did not affect the value of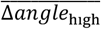 . Taking advantage of this, we generated three cell-type distributions from one distribution: one shifted 12 cells along the X-axis, one shifted 12 cells along the Y-axis, and one shifted 12 cells along both axes.

#### 2. Training a neural network

We used the ResNet18 architecture (30) that was pre-trained with the ImageNet-1K dataset for training. We re-trained the network (i.e., transfer learning) by using Google colaboratory. Hyperparameters used for training were as follows: the optimizer was SGD with a learning rate of 0.01 and a momentum of 0.9. batch size, 50; epoch number, 10. Since ResNet18 accepts images as input, each cell-type distribution was converted to an RGB image with 224 pixels in height and 224 pixels in width each. Since each cell was drawn as a hexagon, the converted image had margins at edges. The red channel was assigned to high cells, the green to low cells, and the blue to the margin. Then intensities of images were normalized so that the mean and SDs of intensities of each channel matched with these of the ImageNet-1K dataset. The accuracy of the trained network was evaluated by counting the number of validation data whose labels were correctly predicted by the network.

### 3. Grad-CAM visualization of the neural network attention

Grad-CAM is an explainable AI technique that can visualize which regions of images the network focuses on when making a classification of an input image (31). We used pytorch-gradcam package in Python to visualize the network attention. Layer2[-1] of PyTorch ResNet18 were selected as a target layer to compute the attention map.

### Generalized linear model

All following procedures were performed in R. Generalized linear models (GLM) were fit to data by using *glm* function of *stats* package (version 4.4.1). Explanatory variables were standardized before fitting GLM by using *scale* function of *base* package (version 4.4.1), which enabled comparisons of coefficients of explanatory variables. Gamma distribution was selected as a probability distribution, because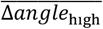 takes positive continuous values. The object variable was related to explanatory variables via a log link function. To avoid the multicollinearity, we calculated variance inflation factors (VIF) by using *vif* function of *car* package (version 3.1-2). We selected explanatory variables so that the value of VIF does not to exceed 10. Then, models were evaluated based on values of AIC (Akaike information criterion) by using *dredge* function of *MuMIN* package (version 1.48.4). The model that took the lowest value of AIC was selected and used in the manuscript. We used *r2* function of package *performance* version 0.12.4 for calculating Pearson correlation coefficient.

### Quantification of low-cell distributions *in vivo*

In Fig. 6A-B’, data adopted from (11) were reanalyzed. In Fig. S7, data adopted from (9) were reanalyzed. Cluster lengths were measured as follows: First, cells were segmented using Cellpose2 (38), and the segmented data were saved as ImageJ ROIs. ROIs marking SCCs were selected based on signals of MCC markers (γ-TUBULIN or acetylated-TUBULIN). These ROIs were then dilated and filled with white, resulting in fusions of areas of adjacent SCCs. The images were then rotated so that the ovary-uterus axis aligned with the X-axis of the image. In the rotated images, white areas were converted to ROIs using the analyze particle function in Fiji. Using these ROIs, the O-U and circumferential lengths of each cluster were measured by the measurement function in Fiji. Clusters that crossed the image edges or epithelial edges were excluded from our analysis. Since oviduct epithelial cells elongate along the ovary-uterus axis (9), we normalized the O-U and circumferential lengths of each SCC cluster. To achieve this, we divided cluster lengths by the average length of SCCs that comprise each cluster. These values were plotted in Fig. 6B and B’’’ and Fig. S7C-D.

In Fig. S7, frequencies of MCCs were measured by calculating the ratio of MCCs to all cells in each image. To measure the frequency of edges between SCCs and MCCs, we counted the number of these edges and divided the value by the number of all cell edges in each image.

### Imaging fluorescent signals

Oviducts were dissected and opened longitudinally along the ovary uterus axis. For imaging VANGL2:EGFP signals, oviducts were fixed in 4% PFA in PBS (-) at 4°c overnight. After washing, samples were incubated with Blocking One (Nacalai Tesque, 03953-95) at room temperature for 30 min. The samples were then incubated with anti-acetylated-TUBULIN antibody (1/500, Sigma-Aldrich, T7451) in Blocking One (Nacalai Tesque, 03953-95) at 4°c overnight. After washing, samples were incubated with phalloidin (1/500, Thermo Fisher Scientific, A12381) and anti-Mouse IgG antibody (1/500, Thermo Fisher Scientific, A21235) in Blocking One (Nacalai Tesque, 03953-95) at 4°c overnight. Finally, after washing, samples were mounted in Fluoromount G (SouthernBiotech, 0100-01). All washing steps were performed as follows: 3 x rinse and 3 x 10 min washes with 0.1% Triton X-100 in PBS (-) at room temperature. Images were acquired using Nikon ECLIPSE Ti2 microscope equipped with CSUW1 spinning disk confocal unit (Yokogawa).

### Quantification of VANGL2:EGFP distributions

In Figure 6, the distribution of VANGL2:EGFP at cell edges of individual cells was quantitatively evaluated using QuantifyPolarity v2.1 (39). The principal component analysis (PCA) method provided by QuantifyPolarity was used to determine both the magnitude and angle of polarity in the VANGL2:EGFP distribution. The angle represents the orientation of the signal-enriched region. The magnitude reflects the degree of directional bias in the signal distribution; higher values (depicted as longer bars in Fig. 6) indicate stronger enrichment in a specific direction along the cell periphery.

### Mouse

Female mice (*R26-VANGL2-EGFP* transgenic mice, (10)) were used to study VANGL2:EGFP distributions in wild-type oviducts. Details of *R26-VANGL2-EGFP* transgenic mice (10) were described in these papers. Animal care and experiments were conducted following the Guidelines of Animal Experimentation of National Institutes for Natural Sciences. All animal experiments were approved by either the Animal Research Committee of National Institutes for Natural Sciences.

Animals were maintained in a light- and temperature-controlled room using a 12 h:12 h light:dark cycle at 23±2°C.

## Acknowledgments

We thank Dr. Goshi Ogita for discussion; members of the T.F laboratory for discussion and technical assistance. This work was supported by JSPS KAKENHI Grant Numbers, 21H02494, 24K02039, 22H05168 (to T.F.), 22K15130 and the Takeda Science Foundation (to M.A.).

## Author Contributions

M.A., H.K. and T. F. designed research; M.A. performed research; M.A. analyzed data; and M.A., H.K. and T. F. wrote the paper.

## Use of generative AI and AI-assisted technologies

Gemini (Google) was used for writing R, Fiji, and Python scripts. ChatGPT (Open AI) was used for improving language and readability of the manuscript. After using these tools, we reviewed and edited the content of the scripts and the manuscript.

