## Supporting Information for "Directional alignment of different cell types organizes planar cell polarity"

**This PDF file includes:**

Supporting text
Figures S1 to S7
Tables S1 to S2
SI References

**Supporting Information Text**

**Methods**

**The mathematical model**

In our mathematical model, epithelial tissue is represented by an array of regular hexagonal cells. In each hexagonal cell, the FZD and VANGL complexes are distributed on the cell edges and within the intracellular region. Temporal changes in the concentration of the FZD complex are described by ordinary differential equations;

$$\frac{dF_{\mathrm{C}_k}}{dt} = -\sum_i K_{\mathrm{F}_{ki}} F_{\mathrm{C}_k} + \sum_i R_{\mathrm{F}_{ki}} F_{\mathrm{M}_{ki}} \qquad (s1)$$

$$\frac{dF_{\mathrm{M}_{ki}}}{dt} = -R_{\mathrm{F}_{ki}} F_{\mathrm{M}_{ki}} + K_{\mathrm{F}_{ki}} F_{\mathrm{C}_k} \qquad (s2)$$

where $F_{\mathrm{C}_k}$ and $F_{\mathrm{M}_{ki}}$ represent concentrations of the FZD complex in the intracellular region and on cell edge $i$ of the $k$th cell, respectively (Fig. 1B). The FZD complex in the intracellular region is transported to cell edge $i$ at a rate of $K_{\mathrm{F}_{ki}}$, while the FZD complex on cell edge $i$ is internalized into the intracellular region at a rate of $R_{\mathrm{F}_{ki}}$ (Fig. 1B, blue arrows). Similarly, temporal changes in the concentration of the VANGL complex are described as follows;

$$\frac{dV_{\mathrm{C}_k}}{dt} = -\sum_i K_{\mathrm{V}_{ki}} V_{\mathrm{C}_k} + \sum_i R_{\mathrm{V}_{ki}} V_{\mathrm{M}_{ki}} \qquad (s3)$$

$$\frac{dV_{\mathrm{M}_{ki}}}{dt} = -R_{\mathrm{V}_{ki}} V_{\mathrm{M}_{ki}} + K_{\mathrm{V}_{ki}} V_{\mathrm{C}_k} \qquad (s4)$$

where $V_{\mathrm{C}_k}$ and $V_{\mathrm{M}_{ki}}$ represent concentrations of the VANGL complex in the intracellular region and on cell edge $i$ of the $k$th cell, respectively (Fig. 1B). The VANGL complex in the intracellular region is transported to the cell edge $i$ at a rate of $K_{\mathrm{V}_{ki}}$, and the VANGL complex on cell edge $i$ is internalized into the intracellular region at a rate of $R_{\mathrm{V}_{ki}}$ (Fig. 1B, blue arrows).To implement autoactivation, mutual inhibition, and intercellular coupling (Fig 1A'), the shuttling rates are described as follows;

$$K_{\mathrm{F}_{ki}} = \rho_{\mathrm{F}} + \omega_{\mathrm{F}} F_{\mathrm{M}_{ki}} V'_{\mathrm{M}_{ki}} \qquad (s5)$$

$$R_{\mathrm{F}_{ki}} = \mu_{\mathrm{F}} + \varphi_{\mathrm{F}} V_{\mathrm{M}_{ki}} F'_{\mathrm{M}_{ki}} \qquad (s6)$$

$$K_{\mathrm{V}_{ki}} = \rho_{\mathrm{V}} + \omega_{\mathrm{V}} V_{\mathrm{M}_{ki}} F'_{\mathrm{M}_{ki}} \qquad (s7)$$

$$R_{\mathrm{V}_{ki}} = \mu_{\mathrm{V}} + \varphi_{\mathrm{V}} F_{\mathrm{M}_{ki}} V'_{\mathrm{M}_{ki}} \qquad (s8)$$

where $F'_{\mathrm{M}_{ki}}$ and $V'_{\mathrm{M}_{ki}}$represents concentrations of the FZD and VANGL complexes on the cell edge of an adjacent cell juxtaposed to cell edge $i$ of the $k$th cell (Fig. 1B). Intercellular bridge formation was implemented by assuming that the autoactivation and mutual inhibition activities of the FZD and VANGL complexes depend on the formation of intercellular FZD-VANGL complex (FZD-VANGL bridge, Fig. 1A'). The concentration of the FZD-VANGL bridge on cell edge $i$ in $k$th cell is assumed to be proportional to $F_{\mathrm{M}_{ki}} V'_{\mathrm{M}_{ki}}$ (or $V_{\mathrm{M}_{ki}} F'_{\mathrm{M}_{ki}}$). This assumption, along with the mass-action terms, is adopted from a previous study (1). The second term of eq. s5 represents the autoactivation activity of the FZD-VANGL bridge. Note that increases in $F_{\mathrm{M}_{ki}}$ and $V'_{\mathrm{M}_{ki}}$on cell edge $i$ elevate the value of $K_{\mathrm{F}_{ki}}$, a transportation rate of the FZD complex from the intracellular region to cell edge $i$. Similarly, the second term of eq. s6 represents the mutual inhibition activity of the FZD-VANGL bridge, where increases in $V_{\mathrm{M}_{ki}}$ and $F'_{\mathrm{M}_{ki}}$on cell edge $i$ elevate the value of $R_{\mathrm{F}_{ki}}$. $\omega$ and $\varphi$ are constants that regulate the strength of autoactivation and mutual inhibition activities, respectively. $\rho$ is a constant that represents the basal shuttling rate of the FZD and VANGL complex from the intracellular region to a cell edge, while $\mu$ is a constant that represents the basal internalization rate of the FZD and VANGL complex from a cell edge to the intracellular region.

Furthermore, eq. s5-8 were modulated as follows to incorporate non-local effects.

$$K_{\mathrm{F}_i} = \rho_F + \omega_F \left( F_{\mathrm{M}_i} V'_{\mathrm{M}_i} + d_{\mathrm{A_F}} \sum_{j=i-1,i+1} F_{\mathrm{M}_j} V'_{\mathrm{M}_j} \right) \tag{s9}$$

$$R_{\mathrm{F}_i} = \mu_F + \varphi_F \left( V_{\mathrm{M}_i} F'_{\mathrm{M}_i} + d_{\mathrm{I_V}} \sum_{j=i-1,i+1} V_{\mathrm{M}_j} F'_{\mathrm{M}_j} \right) \tag{s10}$$

$$K_{\mathrm{V}_i} = \rho_V + \omega_V \left( V_{\mathrm{M}_i} F'_{\mathrm{M}_i} + d_{\mathrm{A_V}} \sum_{j=i-1,i+1} V_{\mathrm{M}_j} F'_{\mathrm{M}_j} \right) \tag{s11}$$

$$R_{V_i} = \mu_V + \varphi_V \left( F_{\mathrm{M}_i} V'_{\mathrm{M}_i} + d_{\mathrm{I_F}} \sum_{j=i-1,i+1} F_{\mathrm{M}_j} V'_{\mathrm{M}_j} \right) \tag{s12}$$

where the second term in parentheses denotes the non-local effects. For example, in eq. s9, an increase in the concentration of the FZD-VANGL bridges on cell edge $i-1$ and $i+1$ elevate the value of $K_{\mathrm{F}_{ki}}$, a transportation rate of the FZD complex from the intracellular region to cell edge $i$ (Fig S2A and A'). $d_{\mathrm{A}}$ and $d_{\mathrm{I}}$ are constants that regulate the strength of non-local effects (Fig S2A and A'). In previous theoretical studies, these non-local effects are implemented either by diffusion of molecules on cell membrane and/or within the intracellular region (1), or by simply assuming that core protein complexes on cell membrane exert autoactivation and/or mutual inhibition effects on neighboring membrane compartments (2). We adopted the latter approach, as the details of non-local effects remain unclear and our study did not require a focus on these details.

Simulations were performed by the Euler's method with a time step $\Delta t = 0.01$. The calculations were iterated until the simulation converged.

The parameter values of the model (listed in Table S1) were optimized so that the model reproduces the non-autonomous effects of mutant cells lacking core proteins on surrounding wild-type cells, as observed *in vivo* (3, 4). Since the non-autonomous effects in the mouse oviduct have not been thoroughly assessed, we referred to findings from *Drosophila* wing PCP, where the non-autonomous effects have been extensively investigated. Each wing epithelial cell produces a wing hair that points to the distal end of the wing. When cells lacking Frizzled (Fz) or Van Gogh (Vang) are genetically produced in the *Drosophila* wing, polarities of surrounding wild-type cells are reorganized (3). In the presence of *fz* mutant cell clusters, wing hair orientations of wild type cells is reversed on the distal side of the cluster (3). In our model, this phenotype was reproduced by introducing a cluster of cells completely lacking the FZD complex (Fig. S3A1). In contrast, near a *vang* mutant cell cluster, the wing hair orientations are reversed on the proximal side (3), a phenotype we replicated by implementing a cluster of cells entirely devoid of the VANGL complex (Fig. S3A2). When *flamingo,* a fly homolog of *Celsrs*, was lost in a cluster of cells, no reversal of wing hair orientation was observed in surrounding wild-type cells. However, when clusters of wild-type cells were surrounded by *flamingo* mutant cells, the wing hairs in the wild-type island formed a swirling pattern (3). Since Flamingo is a component of both the FZD and VANGL complexes, its loss can be approximated by the loss of both complexes. In our model, when cells lacking both FZD and VANGL complexes were introduced, misorientation of polarity was emerged only in the vicinity of mutant cells and no polarity reversal was observed (Fig. S3A3). However, when a cluster of wild-type cells was surrounded by cells lacking both FZD and VANGL complexes, a swirling pattern of polarity coordination emerged (Fig. S3A4), although the emergence of this swirling pattern appeared to depend on the initial distribution of the FZD and VANGL complexes.

**The initial state of the simulation**

We considered two types of hexagonal cell geometries, which have different initial distribution of the FZD and VANGL complexes as shown in Fig. S2B and B' (the initial state #1 and #2). The initial concentration in the intracellular region of $k$ th cell was determined as 2-$\sum_i F_{\mathrm{M}_{ki}}$. $balance_{\mathrm{LH}}$ is a constant that governs the magnitude of the core protein imbalance between high cells and low cells. In low cells, the initial concentrations of the FZD and VANGL complexes at cell edges and in intercellular regions were multiplied by $balance_{\mathrm{LH}}$, thereby reducing their levels accordingly. To manage floating point errors, small random noises were added to the initial concentrations of the FZD and VANGL complexes. The value of noise ranged from 0 to $0.6*10^{-10}$ in high cells and 0 to $balance_{\mathrm{LH}}*0.6*10^{-10}$ in low cells.

In Fig. 5, clusters of low cells were shifted randomly along the X and Y axis of the epithelial sheet. Length of shift, $\sigma$, of each cluster along each axis was determined randomly as $\sigma = floor\left(rand * (\sigma_{max} * 2 + 1)\right) - \sigma_{max}$, where $rand$ is a random number that ranges from $0 < rand < 1$, and $floor(x)$ returns the largest integer not greater than $x$. $\sigma_{max}$ determines the max length of the shift.

**Polarity vectors**

The polarity of the FZD-complex distribution along cell edges was represented by a bar centered within each cell. The angle and the length of the bar represents the orientation of the FZD complex-enriched cell edges and the magnitude of the polarity, respectively. The orientation and the magnitude of polarity vectors were calculated as follows:

$$F_{\mathrm{x}_k} = \sum_{i=1}^{6} F_{\mathrm{M}_{ki}} \cos\theta_{ki}$$

$$F_{\mathrm{y}_k} = \sum_{i=1}^{6} F_{\mathrm{M}_{ki}} \sin\theta_{ki}$$

$$angle_k = \arctan(F_{\mathrm{y}_k}/F_{\mathrm{x}_k})$$

$$magnitude_k = \frac{\sqrt{{F_{\mathrm{x}_k}}^2 + {F_{\mathrm{y}_k}}^2}}{\sum_{i=1}^{6} F_{\mathrm{M}_{ki}}}$$

Here, $angle_k$ and $magnitude_k$ represent the orientation of the FZD complex-enriched cell edges and the magnitude of the polarity of $k$th cell, respectively. $angle_k$ ranges from -180 to 180 degrees. $\theta_{ki}$ is an angle of a line connecting the centroid of cell edge $i$ and the centroid of $k$th cell relative to the X-axis of the cell sheet. For example, $\theta_{ki} = 30$ degrees in Fig. 1B.

**Quantification of the magnitude of PCP disorder**

We quantified the distribution of the FZD complex on cell edges to evaluate the magnitude of the disorder in PCP. Since the equations describing the dynamics of the FZD complex are equivalent to those of the VANGL complex in our model, there is no essential difference whether we use the distributions of the FZD or VANGL complex to evaluate PCP disorder.

To evaluate the magnitude of PCP disorder, we calculated $\overline{\Delta angle_{\mathrm{high}}}$ as follows:

$$\overline{\Delta angle_{\mathrm{high}}} = \frac{1}{n_{\mathrm{high}}} \sum_{m \in \{\text{high cells}\}} \left| angle_{m,\mathrm{initial}} - angle_{m,\mathrm{final}} \right|$$

$n_{\mathrm{high}}$ is a number of high cells in a cell sheet. $angle_{m,\mathrm{initial}}$ and $angle_{m,\mathrm{final}}$ are the angle of the polarity vector of $m$th high cell in a cell sheet at the initial and final state, respectively. Additionally, we calculated a circular variance, which represents the angular distribution of the polarity vectors, to quantify the magnitude of PCP disorder. The circular variance ranges from 0 to 1, where a value of 0 indicates that polarity vectors are unidirectionally oriented, while a value of 1 indicates that vectors are randomly oriented. We preferred $\overline{\Delta angle_{\mathrm{high}}}$, as the circular variance undervalues the disorder of PCP when polarity vectors were misoriented but still unidirectionally aligned.

**Statistics**

The statistical methods used in each experiment are described in the figure legends or within the manuscript. We used R packages as follows. Wilcoxon signed-rank test: *wilcox.test* function of package *stats* version 4.4.1; Wilcoxon rank-sum test: *wilcox.test* function of package *stats* version 4.4.1; Steel-Dwass test: *pSDCFlig* function of package *NSM3* version 1.18

**GLM**

For the GLM analysis in Figure 5 and S5, we generated data with varying cluster intervals to minimize correlations between cluster lengths and $freq_{\mathrm{low}}$. A complete list of parameter values used for data generation is provided in Table S2.

We excluded data simulated with $balance_{\mathrm{LH}}$>0.5 because, in this range, $\overline{\Delta angle_{\mathrm{high}}}$ were only weakly affected by low-cell distributions. Including these data in GLM modeling reduced the predictive accuracy of the model.

**Quantification of CELSR1 immunofluorescent signals and VANGL2:EGFP signals**

For quantification of CELSR1 immunofluorescent signals, data from (5) were reanalyzed. Before quantification, the intensity of background signals was subtracted from the images. To measure background signal intensity in each image, three regions of interest (ROIs), each measuring 30 pixels in height and 30 pixels in width, were placed in areas outside the epithelial fold (Fig. S6A). The average intensity across these ROIs was defined as the intensity of background signals.

Cell edges were segmented using Cellpose2 (6) and skeletonized using the Skeletonize function in Fiji. Each cell-edge was then saved as a ROI (cell-edge ROI) using a Fiji macro. Within each ROI, the average intensity of signals was measured using the measurement function in Fiji, and the CELSR1 or VANGL2:EGFP intensity was normalized by divided it by the F-actin intensity. Cell edges were categorized as either circumferential or O-U based on the orientation of each ROI's centroid relative to the centroid of the corresponding cell (Fig. S6B). Additionally, cell edges were further classified into three groups: edges between two MCCs (MCC-MCC), edges between as MCC and as SCC (MCC-SCC), and edges between two SCCs (SCC-SCC). Cell types (MCC or SCC) were determined based on acetylated-TUBULIN immunofluorescent signals.

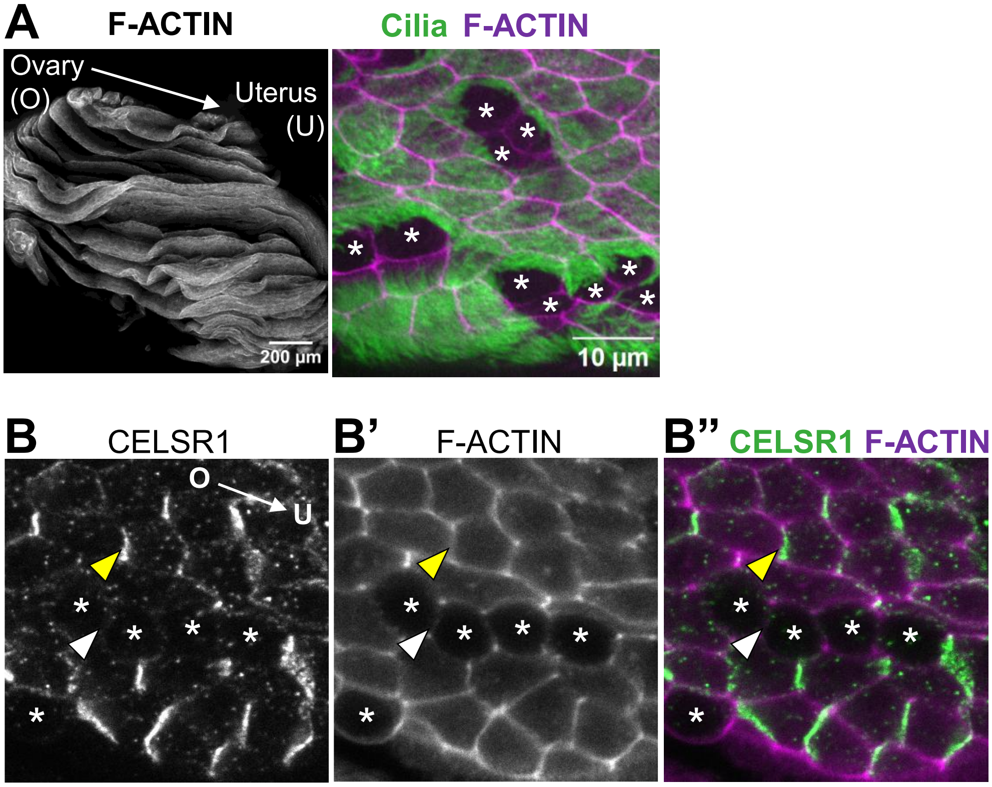


**Figure S1.** Core protein imbalance in the mouse oviduct epithelium
(A) The infundibulum of the oviduct was dissected, longitudinally opened along the ovary (O)-uterus (U) axis, and stained for F-actin (gray in the left panel and magenta in the right panel, a marker for cell edges) and acetylated-TUBULIN (green in the right panel, a marker for cilia). The oviduct lumen is lined with an epithelium composed of MCCs (multiciliated cells) and SCCs (secretory cells). Apical views of the epithelia are shown. (B-B”) Oviduct epithelium stained for CELSR1 (B, green in B”) and F-actin (B’, magenta in B”). CELSR1 is enriched at cell edges perpendicular to the ovary-uterus axis. CELSR1 levels are higher at cell edges between MCCs (yellow arrowhead) than at those between SCCs (white arrowhead). Asterisks: SCCs.

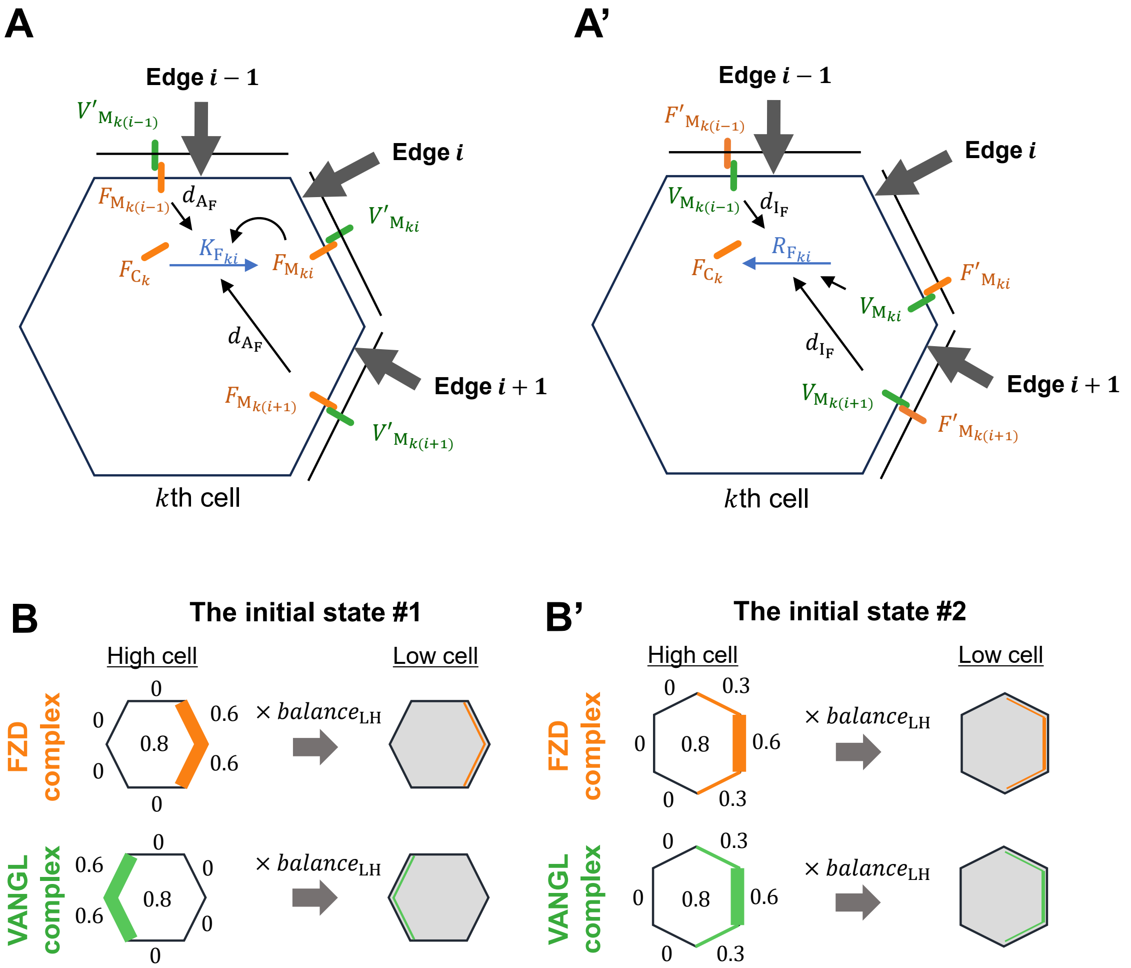


**Figure S2.** Modeling the interactions between core proteins

(**A-A'**) Diagram illustrating the non-local effects of autoactivation (**A**) and mutual inhibition (**A'**) on shuttling rate of the FZD complex. $F_{\mathrm{C}_k}$ and $F_{\mathrm{M}_{ki}}$ represent concentrations of the FZD complex in intracellular region and on cell edge $i$ of the $k$th cell, respectively. The FZD complex in the intracellular region is transported to the cell edge $i$ at a rate $K_{\mathrm{F}_{ki}}$, while the FZD complex on cell edge $i$ is internalized into the intracellular region at a rate $R_{\mathrm{F}_{ki}}$. Similarly, $V_{\mathrm{C}_k}$ and $V_{\mathrm{M}_{ki}}$ represent the concentrations of the VANGL complex in the intracellular region and on cell edge $i$ of the $k$th cell, respectively. $F'_{\mathrm{M}_{ki}}$ and $V'_{\mathrm{M}_{ki}}$ denote the concentrations of the FZD and VANGL complexes on the adjacent cell edge juxtaposed with cell edge $i$ of the $k$th cell, respectively. The rates $K_{\mathrm{F}_{ki}}$ and $R_{\mathrm{F}_{ki}}$ are influenced by the concentration of the FZD and VANGL complexes on cell edge $i-1$, $i$ and $i+1$ (black arrowheads). The strength of effects of the FZD and VANGL complexes on cell edge $i-1$ and $i+1$ on $K_{\mathrm{F}_{ki}}$ and $R_{\mathrm{F}_{ki}}$ is determined by constants $d_{\mathrm{A}}$ and $d_{\mathrm{I}}$, respectively. (**B-B'**) Distribution of the FZD and VANGL complexes in high cells and low cells at the initial state of our simulation. The number at the center of each cell indicates the concentration of the FZD or VANGL complex in the intracellular region, while the number near the cell edge represents a concentration of these complexes on each cell edge. Two types of hexagonal cell geometries were considered in this study. In both geometries, the FZD and VANGL complexes were initially enriched at the right and left ends of the cell, respectively, but the vertex orientations differed.

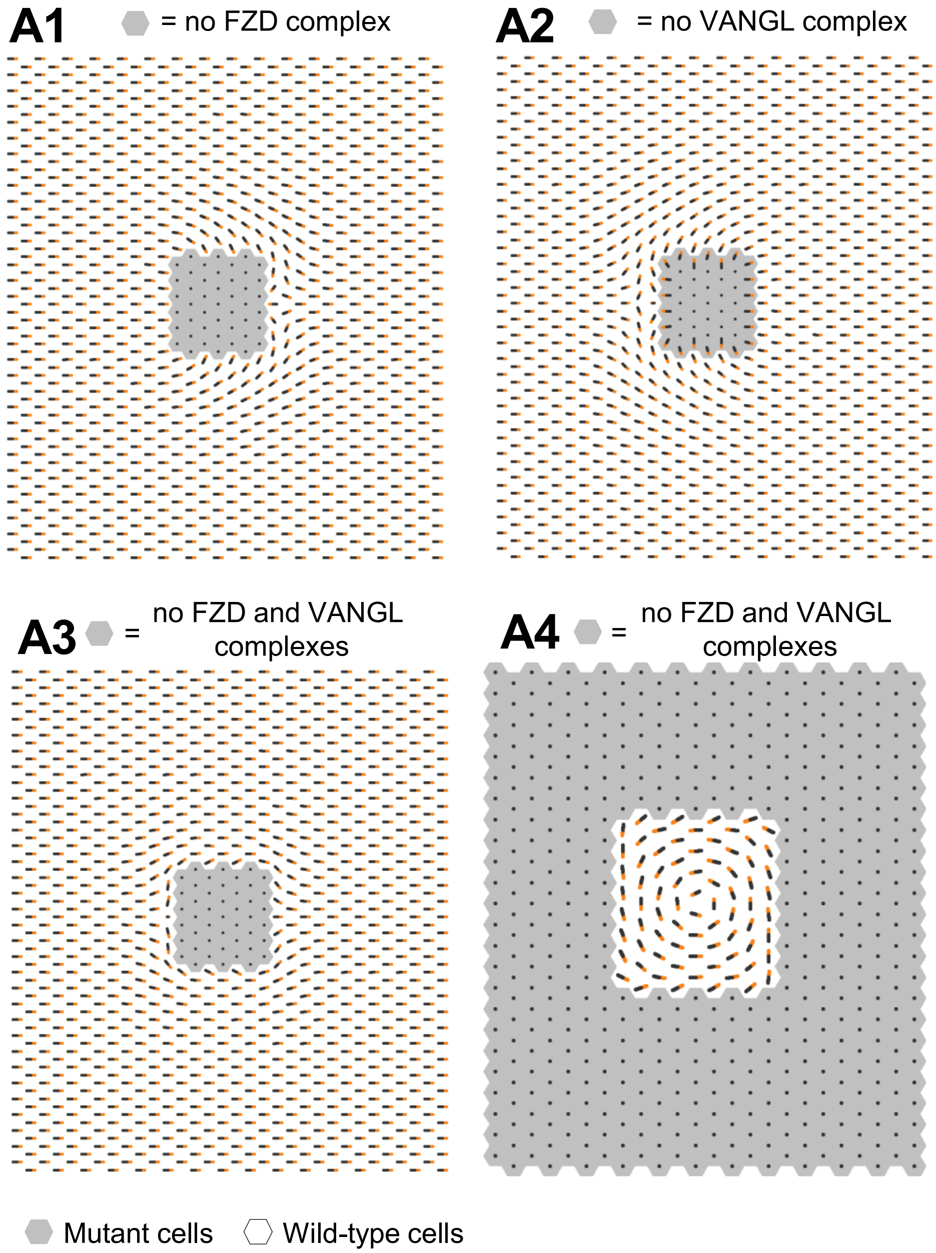


**Figure S3.** Model reproduced non-autonomous effects of cells lacking core proteins
Gray hexagons represent mutant cells completely lacking the FZD complex (**A1**), the VANGL complex (**A2**) or both complexes (**A3** and **A4**). White hexagons represent wild-type cells. The orientations of polarity vectors were reversed on the right side of the FZD-complex-lacking cells (**A1**), and on the left side of the VANGL-complex-lacking cells (**A2**). (**A3-A4**) Effects of mutant cells lacking both the FZD and VANGL complexes on the polarity vectors of surrounding wild-type cells. (**A3**) Polarity vectors were misoriented only in the vicinity of mutant cells and no reversal of polarity vectors was observed. (**A4**) When wild type cells were surrounded by these mutant cells, a swirling pattern of polarity vectors emerged within the island of wild-type cells. In this data, the FZD and VANGL complexes were randomly distributed on cell edges of wild-type cells at the initial timepoint of the simulation.

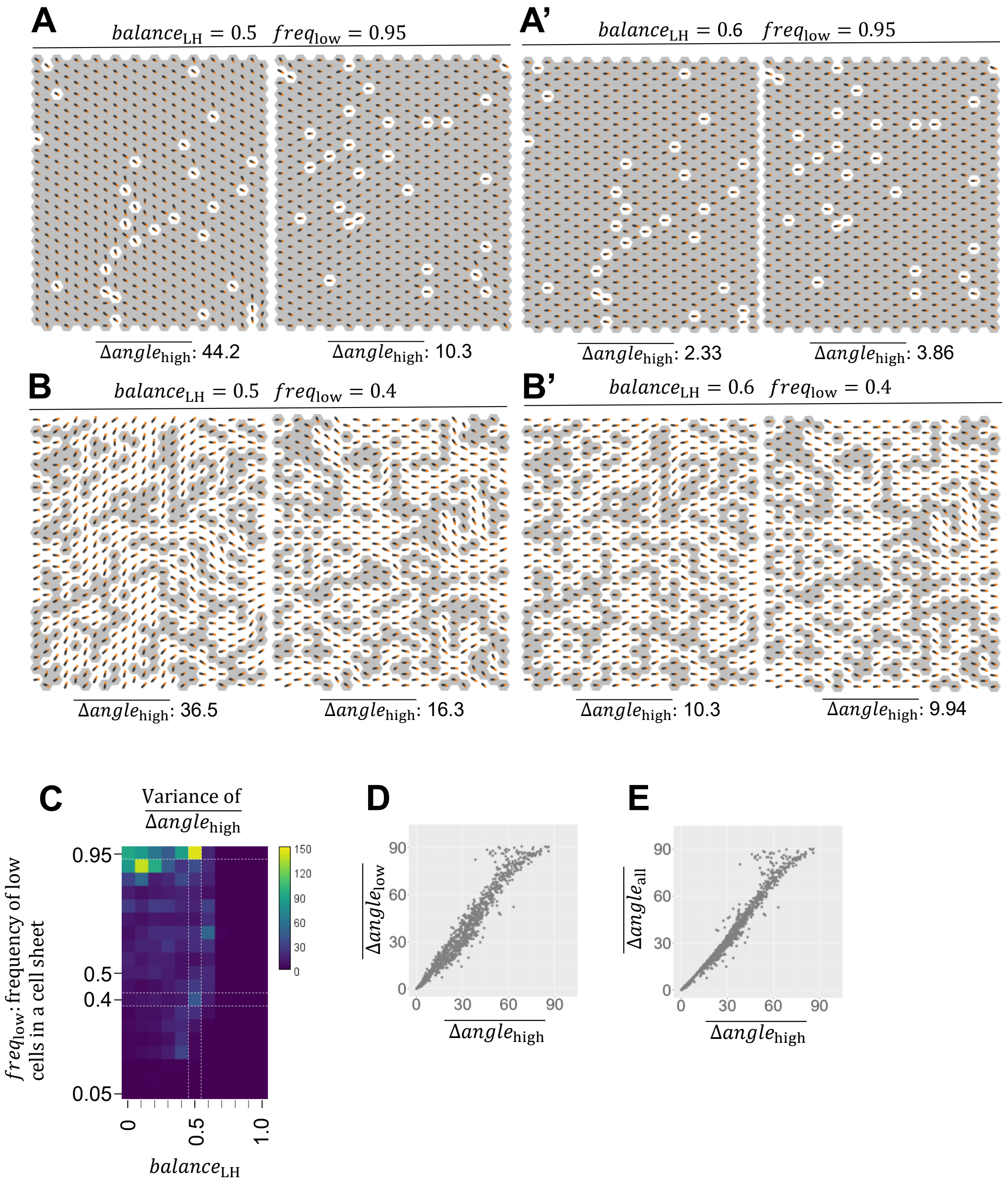


**Figure S4.** Variability in the effects of random low-cell distributions on values of $\overline{\boldsymbol{\Delta angle_{high}}}$

(**A**) Cell-type distributions that resulted in the max and the minimum value of $\overline{\Delta angle_{\mathrm{high}}}$ (left and right, respectively) at $balance_{\mathrm{LH}} = 0.5$ and $freq_{\mathrm{low}} = 0.95$ in a dataset shown in Fig. 3A-B. Effects of these cell-type distributions on PCP at $balance_{\mathrm{LH}} = 0.6$ are shown in (**A'**). (**B**) Cell-type distributions that took the max and the minimum value of $\overline{\Delta angle_{\mathrm{high}}}$ (left and right, respectively) at $balance_{\mathrm{LH}} = 0.5$ and $freq_{\mathrm{low}} = 0.4$. Effects of these cell-type distributions on PCP at $balance_{\mathrm{LH}} = 0.6$ are shown in (**B'**). (**C**) Heat map showing variance in values of $\overline{\Delta angle_{\mathrm{high}}}$ of 10 random distributions at each set of parameter values of $balance_{\mathrm{LH}}$ and $freq_{\mathrm{low}}$. (**D** and **E**) Scatter plots showing correlation between $\overline{\Delta angle_{\mathrm{high}}}$ and $\overline{\Delta angle_{\mathrm{low}}}$ (**D**) and between $\overline{\Delta angle_{\mathrm{high}}}$ and $\overline{\Delta angle_{\mathrm{all}}}$ (**E**). Each dot represents one simulation.

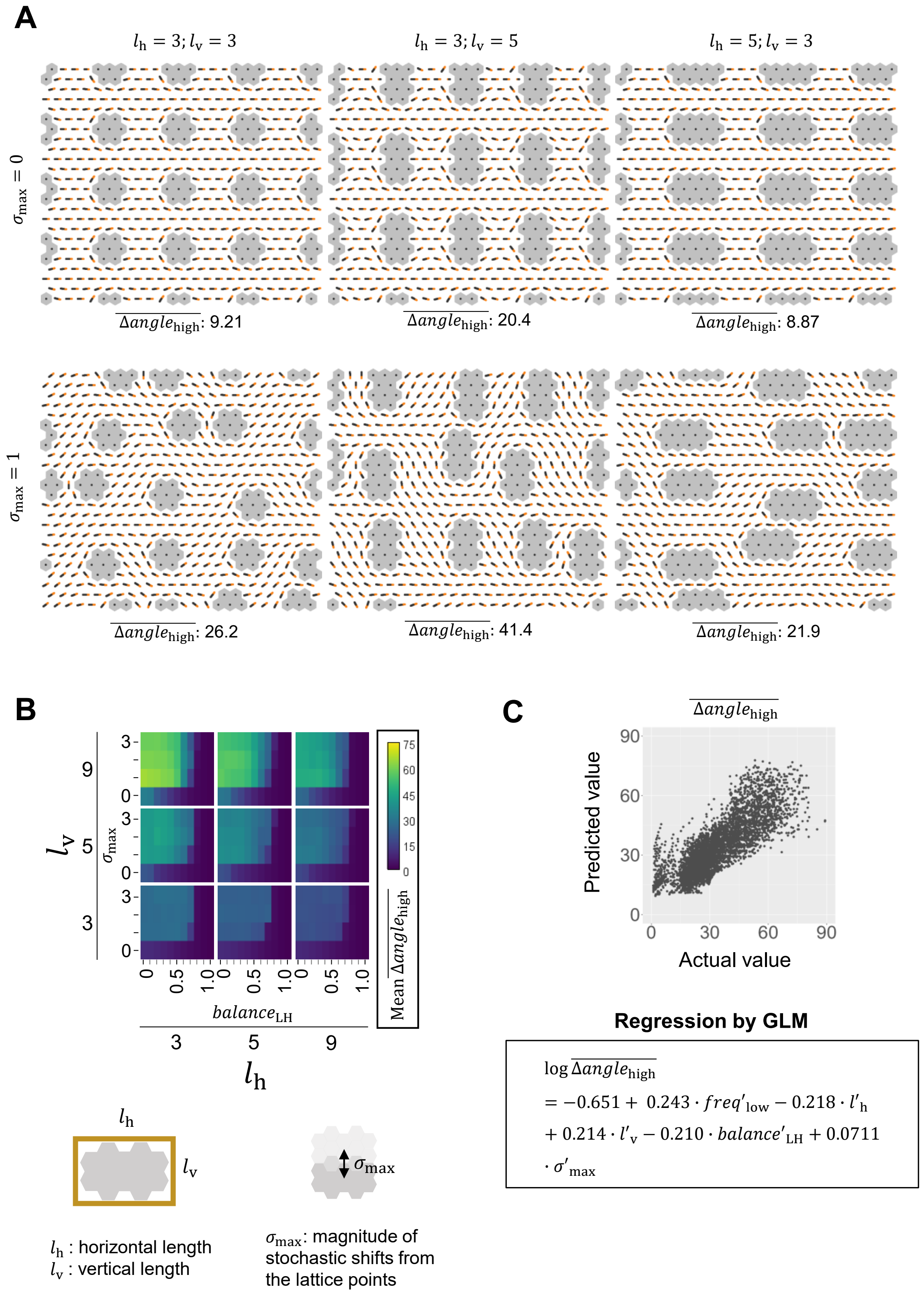
A
$l_h = 3; l_v = 3$
$l_h = 3; l_v = 5$
$l_h = 5; l_v = 3$
$\sigma_{max} = 0$
$\sigma_{max} = 1$
$\overline{\Delta angle_{high}}$: 9.21
$\overline{\Delta angle_{high}}$: 20.4
$\overline{\Delta angle_{high}}$: 8.87
$\overline{\Delta angle_{high}}$: 26.2
$\overline{\Delta angle_{high}}$: 41.4
$\overline{\Delta angle_{high}}$: 21.9
B
$l_v$
$\sigma_{max}$
$balance_{LH}$
$l_h$
Mean $\overline{\Delta angle_{high}}$
$l_h$ : horizontal length
$l_v$ : vertical length
$\sigma_{max}$: magnitude of stochastic shifts from the lattice points
C
$\overline{\Delta angle_{high}}$
Predicted value
Actual value
Regression by GLM
$\log \overline{\Delta angle_{high}} = -0.651 + 0.243 \cdot freq'_{low} - 0.218 \cdot l'_h + 0.214 \cdot l'_v - 0.210 \cdot balance'_{LH} + 0.0711 \cdot \sigma'_{max}$

**Figure. S5.** Effects of low-cell cluster lengths are not affected by hexagonal cell geometries

The geometry shown in Figure S2B' is used in this analysis. (**A**) $l_{\mathrm{h}}$: length of cluster along the X-axis of the cell sheet; $l_{\mathrm{v}}$: length of cluster along the Y-axis of the cell sheet; Clusters were aligned in a lattice (top panels) and were randomly shifted $\sigma_{\mathrm{max}}$ cell(s) at maximum along X and Y axis of each image (bottom panels). (**B**) Heatmaps showing mean values of $\overline{\Delta angle_{\mathrm{high}}}$, averaged over 10 distributions for each set of $balance_{\mathrm{LH}}$, $\sigma_{\mathrm{max}}$, $l_{\mathrm{h}}$, and $l_{\mathrm{v}}$. Colors represent mean $\overline{\Delta angle_{\mathrm{high}}}$ values as shown in the right lookup table (unit: degrees). (**C**) Scatter plot showing the correlation between actual $\overline{\Delta angle_{\mathrm{high}}}$ values and those predicted by the GLM (bottom panel). Pearson correlation coefficient: 0.818. Each dot represents a single simulation.

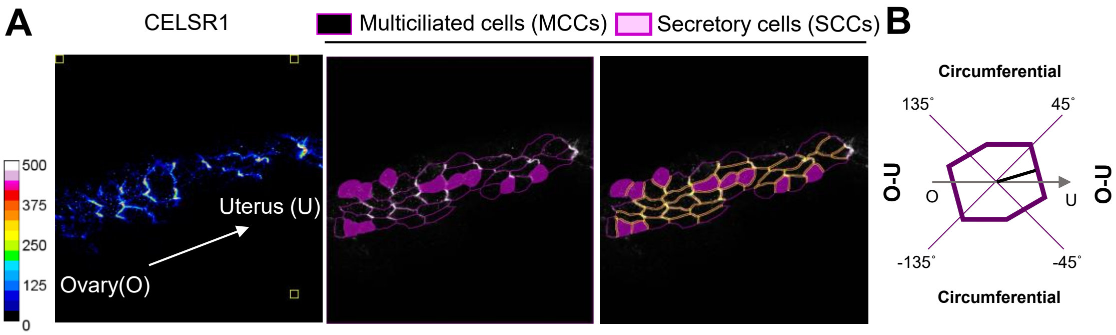


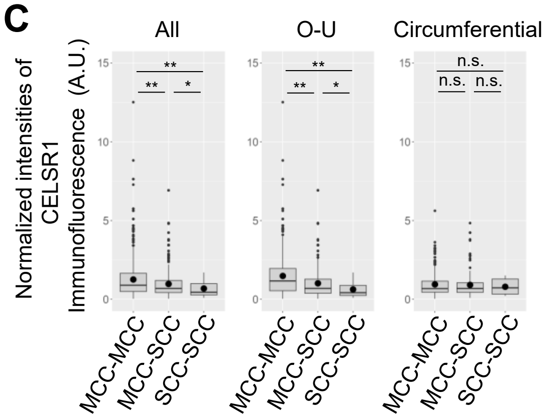


C'

| | Edge | Mean (SD) | N (edges) |
|---|---|---|---|
| All | MCC-MCC | 1.26 (1.19) | 815 |
| | MCC-SCC | 0.973 (0.925) | 464 |
| | SCC-SCC | 0.685 (0.502) | 72 |
| O-U | MCC-MCC | 1.48 (1.36) | 474 |
| | MCC-SCC | 1.01 (1.00) | 275 |
| | SCC-SCC | 0.627 (0.519) | 47 |
| Circum-ferential | MCC-MCC | 0.945 (0.787) | 341 |
| | MCC-SCC | 0.912 (0.798) | 189 |
| | SCC-SCC | 0.794 (0.457) | 25 |

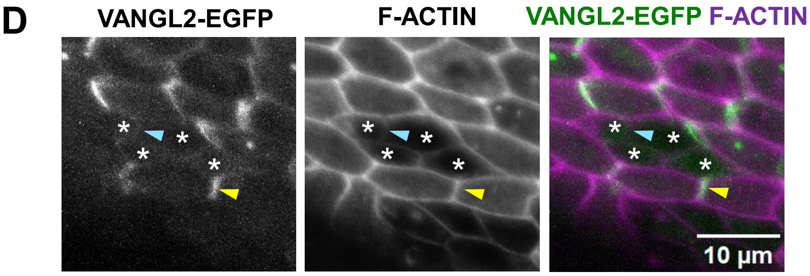


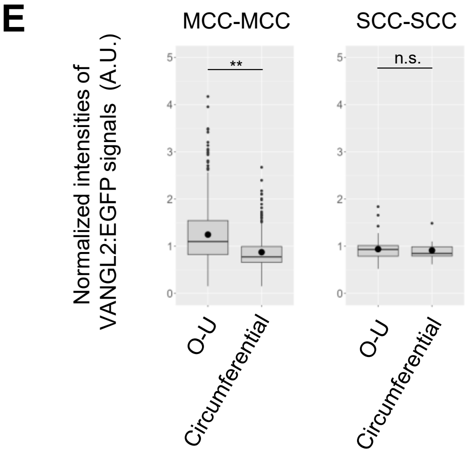


E'

| | Edge | Mean (SD) | N (edges) |
|---|---|---|---|
| MCC-MCC | O-U | 1.25 (0.606) | 1010 |
| | Circumferential | 0.873 (0.343) | 620 |
| SCC-SCC | O-U | 0.938 (0.271) | 79 |
| | Circumferential | 0.912(0.218) | 25 |

**Figure S6.** Oviduct secretory cells are low cells

(**A-C'**) Quantification of CELSR1 immunofluorescence intensities. (**A**) (left) Intensities of CELSR1 immunofluorescent signals are shown by a heat map. Three 30 × 30-pixel yellow boxes were placed outside the epithelial sheet in each image; the mean intensity within these boxes was taken as background. (mid) SCCs are filled with magenta. Magenta lines represent positions of cell edges. CELSR1 signals are shown in white. (right) Each cell edge defined by a pair of vertices was segmented and outlined as a yellow region of interest (ROI). (**B**) Diagram illustrating the categorization of cell edges based on their direction. A black line connects the centroid of each cell edge and the centroid of a cell. The angle of this line relative to the O-U axis was used to categorize each cell edge as either an O-U or circumferential edge. For example, when the angle was 10 degrees, the edge was categorized as an O-U edge. (**C**) Boxplots showing CELSR1 intensity on cell edges, normalized to F-actin intensity. Edges are grouped by orientation (O–U vs. circumferential) and by cell-pair type (MCC–MCC, MCC–SCC, SCC–SCC). Large black points represent average values. **: p<0.01; *p<005; n.s.: not significant (Steel-Dwass test). (**C'**) Mean and standard deviation (SD) values for each group shown in (**C**). (**D**) The oviduct expressing VANGL2:EGFP (left and green in right panel) under the control of Rosa26 promoter was stained for F-ACTIN (middle and magenta in right panel, a marker for cell edges). Yellow arrowhead: a cell edge between two MCCs. Cyan arrowhead: a cell edge between two SCCs. Asterisks mark SCCs. (**E**)Intensities of VANGL2:EGFP signals were compared between O-U and circumferential edges. Left: a boxplot showing the intensities on edges between two MCCs. Right: a boxplot showing the intensities on edges between two SCCs. **: p<0.01; *p<005; n.s.: not significant (Wilcoxon rank sum test). (**E'**) Mean and standard deviation (SD) values for each group shown in (**E**).

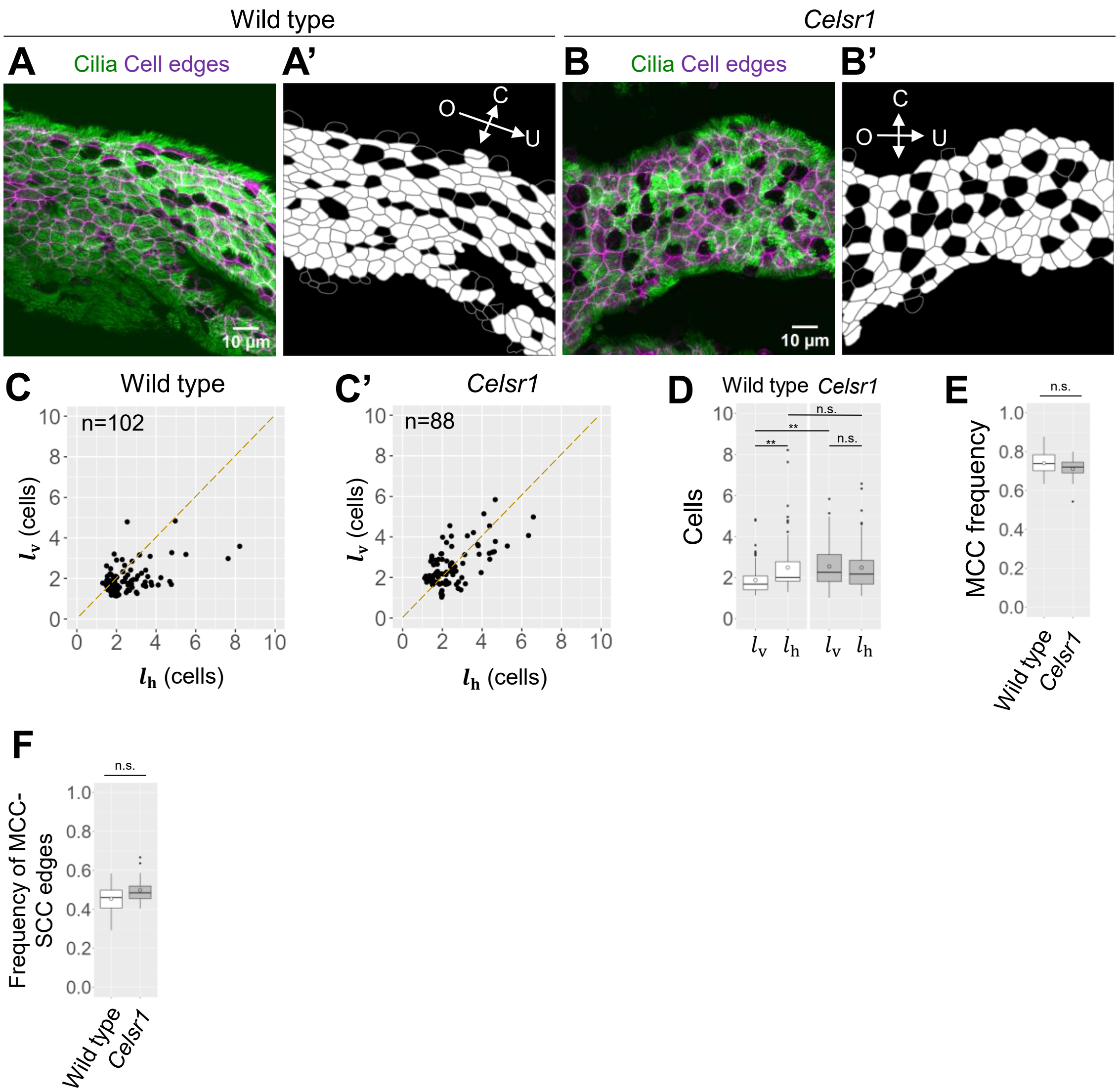


**Figure S7.** CELSR1 aligns SCCs along the ovary-uterus axis

(**A-B**) Oviduct epithelium of 11-week-old mice stained for F-actin (a cell edge marker, magenta) and acetylated-TUBULIN (a cilia marker, green). Images of wild-type (**A-A'**) and *Celsr1* mutant (**B-B'**) tissues are shown. (**A'** and **B'**) Black cells: SCCs; white cells: MCCs; gray lines: cell edges. (**C-D**) $l_h$ and $l_v$ were analyzed by a scatter plot (**C-C'**) and boxplot (**D**). **: p<0.01; n.s.: not significant (Steel-Dwass test). n: number of clusters. Isolated SCCs were excluded from the plots. Each dot represents an SCC cluster. (**E**) MCC frequencies in wild type and *Celsr1* mutant oviducts. n.s., not significant; Wilcoxon rank-sum test; n=20 view fields, 5 wild type oviducts; n=20 view fields, 5 *Celsr1* mutant oviducts. (**F**) Frequencies of edges between MCCs and SCCs in wild type and *Celsr1* mutant oviducts. n.s.: not significant; Wilcoxon rank-sum test. n=20 view fields, 5 wild type mice; n=20 view fields, 5 *Celsr1* mutant oviducts. Open circles in (**D-F**) represent average values.

**Table S1.** List of parameter values used in the mathematical model

| Parameters | Value |
|---|---|
| $\rho_{\mathrm{F}}$ and $\rho_{\mathrm{V}}$ | 0.1 |
| $\mu_{\mathrm{F}}$ and $\mu_{\mathrm{V}}$ | 1 |
| $\omega_{\mathrm{F}}$ and $\omega_{\mathrm{V}}$ | 10 |
| $\varphi_{\mathrm{F}}$ and $\varphi_{\mathrm{V}}$ | 1 |
| $d_{\mathrm{F_A}}$ and $d_{\mathrm{V_A}}$ | 1 |
| $d_{\mathrm{V_I}}$ and $d_{\mathrm{F_I}}$ | 1 |

**Table S2.** List of parameter values used for simulating the effects of low-cell cluster lengths

| $l_h$ | $l_v$ | Interval | $\sigma_{max}$ | $balance_{LH}$ | Related figures |
|---|---|---|---|---|---|
| 3 | 3 | 3 | 0, 1, 2, and 3 | 0-1, interval: 0.1 | Fig 5 and S5 |
| 3 | 5 | 3 | 0, 1, 2, and 3 | 0-1, interval: 0.1 | Fig 5 and S5 |
| 3 | 9 | 3 | 0, 1, 2, and 3 | 0-1, interval: 0.1 | Fig 5 and S5 |
| 5 | 3 | 3 | 0, 1, 2, and 3 | 0-1, interval: 0.1 | Fig 5 and S5 |
| 5 | 5 | 3 | 0, 1, 2, and 3 | 0-1, interval: 0.1 | Fig 5 and S5 |
| 5 | 9 | 3 | 0, 1, 2, and 3 | 0-1, interval: 0.1 | Fig 5 and S5 |
| 9 | 3 | 3 | 0, 1, 2, and 3 | 0-1, interval: 0.1 | Fig 5 and S5 |
| 9 | 5 | 3 | 0, 1, 2, and 3 | 0-1, interval: 0.1 | Fig 5 and S5 |
| 9 | 9 | 3 | 0, 1, 2, and 3 | 0-1, interval: 0.1 | Fig 5 and S5 |
| 3 | 3 | 1 | 0, 1, 2, and 3 | 0-1, interval: 0.1 | Fig 5C and S5C |
| 3 | 5 | 1 | 0, 1, 2, and 3 | 0-1, interval: 0.1 | Fig 5C and S5C |
| 3 | 7 | 1 | 0, 1, 2, and 3 | 0-1, interval: 0.1 | Fig 5C and S5C |
| 5 | 3 | 1 | 0, 1, 2, and 3 | 0-1, interval: 0.1 | Fig 5C and S5C |
| 5 | 5 | 1 | 0, 1, 2, and 3 | 0-1, interval: 0.1 | Fig 5C and S5C |
| 5 | 7 | 1 | 0, 1, 2, and 3 | 0-1, interval: 0.1 | Fig 5C and S5C |
| 7 | 3 | 1 | 0, 1, 2, and 3 | 0-1, interval: 0.1 | Fig 5C and S5C |
| 7 | 5 | 1 | 0, 1, 2, and 3 | 0-1, interval: 0.1 | Fig 5C and S5C |
| 7 | 7 | 1 | 0, 1, 2, and 3 | 0-1, interval: 0.1 | Fig 5C and S5C |
| 3 | 3 | 5 | 0, 1, 2, and 3 | 0-1, interval: 0.1 | Fig 5C and S5C |
| 3 | 3 | 9 | 0, 1, 2, and 3 | 0-1, interval: 0.1 | Fig 5C and S5C |
| 5 | 5 | 7 | 0, 1, 2, and 3 | 0-1, interval: 0.1 | Fig 5C and S5C |
